# A new method based on the von Mises-Fisher distribution shows that a minority of liver-localized CD8 T cells display hard-to-detect attraction to *Plasmodium*-infected hepatocytes

**DOI:** 10.1101/2020.12.12.422451

**Authors:** Viktor S. Zenkov, James O’Connor, Ian Cockburn, Vitaly V. Ganusov

## Abstract

Malaria is a disease caused by *Plasmodium* parasites, resulting in over 200 million infections and 400,000 deaths every year. A critical step of malaria infection is when sporozoites, injected by mosquitoes, travel to the liver and form liver stages. Malaria vaccine candidates which induce large numbers of malaria-specific CD8 T cells in mice are able to eliminate all liver stages, preventing fulminant malaria. However, how CD8 T cells find all parasites in 48 hours of the liver stage lifespan is not well understood. Using intravital microscopy of murine livers, we generated unique data on T cell search for malaria liver stages within a few hours after infection. To detect attraction of T cells to an infection site, we used the von Mises-Fisher distribution in 3D, similar to the 2D von Mises distribution previously used in ecology. Our results suggest that the vast majority (70-95%) of malaria-specific and non-specific liver-localized CD8 T cells did not display attraction towards the infection site, suggesting that the search for malaria liver stages occurs randomly. However, a small fraction (15-20%) displayed weak but detectable attraction towards parasites which already had been surrounded by several T cells. We found that speeds and turning angles correlated with attraction, suggesting that understanding mechanisms that determine the speed of T cell movement in the liver may improve the efficacy of future T cell-based vaccines. Stochastic simulations suggest that a small movement bias towards the parasite dramatically reduces the number of CD8 T cells needed to eliminate all malaria liver stages, but to detect such attraction by individual cells requires data from long imaging experiments which are not currently feasible. Importantly, as far as we know this is the first demonstration of how activated/memory CD8 T cells might search for the pathogen in nonlymphoid tissues a few hours after infection. We have also established a framework for how attraction of individual T cells towards a location in 3D can be rigorously evaluated.

## 1 Introduction

Malaria is a disease caused by parasites of the genus *Plasmodium* that kills over 400,000 people every year^1^. Mosquitoes carrying malaria sporozoites inject sporozoites when searching for blood^2^. Sporozoites travel through the bloodstream to the liver, invade hepatocytes, and form liver stages. The liver stage of malaria infection is asymptomatic. The development of the liver stage takes 48 hours in mice and 7 days in humans^3–6^. Importantly, vaccines inducing exclusively malaria-specific CD8 T cells are capable of removing all liver stages in mice, thus preventing clinical malaria^7–9^. Intravital imaging experiments showed that soon after infection, activated *Plasmodium*-specific CD8 T cells and CD8 T cells of irrelevant specificity cluster around *Plasmodium*-infected hepatocytes^10,11^. Mathematical modeling-based analysis of the T cell clustering data suggested that the formation of the clusters is best explained by a model in which the first T cell finds the liver stage randomly and the attraction of other T cells (including T cells with irrelevant specificity) to the parasite increases with the number of T cells per cluster^11^. While an earlier study suggested that there may be attraction of distant T cells to the clustered liver stage^12^, it remains unclear if attraction to the liver stage occurs prior to the formation of T cell clusters around parasites, as well as if attraction is exhibited by all or just a subset of cells.

Based on intravital imaging experiments, in many if not most analyzed cases, T cells appear to move randomly in tissues *in vivo*^13^. Few studies have accurately quantified if motile T cells exhibit biased migration. In part, this is because of the difficulty in quantifying bias in movement patterns of T cells and relating the movement bias to specific structures in the tissue. Previous studies on the movement of naive B cells in B cell zones of the lymph nodes showed a difference in migration towards the boundary of the follicle, or the plane separating light and dark zones of the germinal center reactions^14,15^. A more rigorous analysis of movement of virus-specific CD8 T cells in the skin demonstrated attraction of T cells to infection sites^16^. However, these studies employed simple metrics to measure attraction (e.g., percent of cells moving to a particular area or the distance between a cell and the boundary) and in most cases required a comparison group to estimate bias. Also, there was no analysis into if any of these metrics have biases. For example, T cells tend to move with a persistent random walk in which a movement’s direction tends to be similar to the previous movement’s direction^17^, which may generate an illusion of a bias towards a specific location. While methods to detect attraction to specific locations have been proposed and studied extensively in ecology^18–20^, ecological movement data are typically 2D. Whether previously proposed methods apply to 3D situations (e.g., for T cells moving in the liver) has not been thoroughly investigated.

At a fundamental level, it is mostly unknown how vaccine-induced T cells search for *in vivo* sites of pathogen replication in peripheral tissues in the hours after infection. This remains an experimental challenge because at early time points, individual pathogens may have too low fluorescent signal to be easily detectable experimentally, and it is difficult to localize memory CD8 T cells near infection sites. To address this fundamental knowledge gap and to understand how CD8 T cells search for the *Plasmodium* liver stages, we performed novel intravital microscopy-based experiments with murine livers in which we tracked positions of liver-localized fluorescently labeled malaria-specific CD8 T cells, CD8 T cells of irrelevant specificity, and malaria liver stages. In our experiments, *Plasmodium* sporozoites expressed GFP at a sufficiently high level so that individual parasites could be followed in the liver minutes to hours after intravenous inoculation^21^.

To evaluate if T cells display attraction towards infection sites, we first utilized previously-used metrics (and developed a correction to an intuitive metric based on the change in distance between T cells and the parasite), then ultimately developed a new metric based on applying the von Mises-Fisher (vMF) distribution to angles to a parasite. This new metric is more powerful (requires less data to detect deviations from random movement) than previously used metrics. Similar to how we found bias in the existing distance metric, we found that our vMF distribution-based metric can be slightly biased for T cells with a correlated/persistent random walk, but we use simulations to approximately correct that bias. Our results suggest that CD8 T cells do not display attraction towards the infection site, with one exception: when a liver stage already possesses a CD8 T cell cluster, a minority of T cells do display strong attraction to this infection site. Stochastic simulations based on the vMF distribution suggest that the detection of weak attraction of individual cells towards the infection site requires amounts of data that are difficult-to-impossible to collect with current intravital imaging protocols. Our work establishes a rigorous framework for evaluating the attraction of moving cells towards a particular location, and begins to explain how malaria-specific CD8 T cells navigate in the presence of malaria infection.

## 2 Materials and Methods

### 2.1 Experimental design

Our data consists of 3D positions over time of CD8 T cells specific for malaria sporozoites; malaria liver stages; and in some cases, control CD8 T cells specific to irrelevant antigens. We analyzed datasets from five sets of experiments: three datasets generated for this analysis and two from a published study^10^. Experimental parameters are described below. For all mice, a lateral incision was made over the left lobe of the liver and the liver was exposed; then the mouse was transfered to the microscope system for imaging. Comprehensive experimental details are provided in the Supplemental information.

For the new **“unclustered/small clustered”** (dataset #1) and **“large clustered”** (dataset #2) datasets, mice were infected with GFP-expressing *Plasmodium berghei* (Pb-CS^5M^) sporozoites, which carry the SIINFEKL epitope from OVA^22,23^. Activated CD8 T cells, specific to SIINFEKL epitope (OT1), were generated *in vitro* by co-culture of naive TCR transgenic CD8 T cells with SIINFEKL peptide as described previously^23^. As a control, we used TCR-transgenic CD8 T cells, specific to GP33 epitope from LCMV (P14) that were activated similarly to the OT1 cells. Our B6 mice received 5 × 10^6^ activated Pb-specific (OT1) or LCMV-specific (P14) CD8 T cells and then 1.5-2 hours later, were infected with 10^5^ Pb-CS^5M^ sporozoites (Figure S1). We performed imaging between 30 min and 2 hours after sporozoite infection using a two-photon microscope^23^. The unclustered/small clustered dataset experiments featured no or few T cells located within 40 *μ*m from the parasite at the beginning of the experiment, and the large clustered dataset experiments featured several T cells located within 40 *μ*m from the parasite (i.e., there was a T cell cluster at the beginning of the experiment^10^). We used a 40 *μ*m radius to distinguish closeness because this value has been used to represent the average radius of a standard murine hepatocyte when roughly modeled as a sphere^11^. The unclustered/small clustered dataset contains 3D coordinates over 3 hours (with timesteps of 1.5 or 2 minutes between recorded stacks of images) of Pb-specific (OT1) and LCMV-specific (P14) CD8 T cells in 4 mice (1 parasite per mouse, Figures S1 and S2). The large clustered dataset contains 3D coordinates over 3 hours (with timesteps of 1 or 2 minutes between recorded stacks of images) of Pb-specific and LCMV-specific CD8 T cells for 3 mice (1 parasite per mouse). As a control, we also performed an experiment in which naive B6 mice received 5 × 10^6^ activated Pb-specific (OT1) cells but no sporozoites (i.e., no infection was given), and livers of the mice were imaged 1.5-2 hours after T cell transfer. This **“No Parasite”** dataset (dataset #3) contains 3D coordinates over 30 minutes (with timesteps of 30 seconds between recorded stacks of images) for 1 mouse.

We also analyzed datasets from our previous study^10^. To generate the **“Paris”** dataset (dataset #4), the following experimental set-up was used. Activated CD8 T cells specific for *Plasmodium yoelii* (Py) sporozoites (PyTCR) were generated *in vivo* by infecting Balb/c mice with the Vaccinia virus expressing the circumsporozoite (CS) protein from Py^10^. Balb/c mice were first infected with Py sporozoites, then 20 hours later, activated Py-specific (PyTCR) CD8 T cells (5 × 10^6^ per mouse) were transferred to the infected mice intravenously. Imaging of the livers of these infected mice was performed 6 hours after the T cell transfer^10^. To generate the **“co-clustered”** dataset (dataset #5), PyTCR cells were activated as described above. In addition, CD8 T cells specific to chicken ovalbumin (OT1) were activated by infecting B6 mice with the Vaccinia virus expressing OVA. CB6 mice (F1 progeny of B6 and Balb/c mice) were infected with 10^5^ Py sporozoites; 20 hours later activated Py-specific (PyTCR, 5 × 10^6^ per mouse) and OVA-specific (OT1, 5 × 10^6^ per mouse) CD8 T cells were transferred into infected mice. Six hours later, livers of these mice were imaged. In both experiments, we performed intravital imaging using spinning-disk confocal microscopy. The Paris dataset contains 3D coordinates over 5 hours (with timesteps of 1, 2, or 4 minutes between recorded stacks of images) of Py-specific CD8 T cells for 26 parasites in 4 mice. The co-clustered dataset contains 3D coordinates over 40 minutes (with timesteps of 2 minutes between recorded stacks of images) of Py-specific and OVA-specific CD8 T cells for 1 parasite in 1 mouse. Note that the terms “parasite” and “infection site” are used interchangeably in much of the paper, as well as “liver stage” when discussing the malaria lifecycle.

All 3D coordinate data are available as a Supplemental information to this paper. We also provide a Mathematica-based script that allows one to estimate the bias of moving agents to a point or a plane based on our newly developed vMF distribution-based metric, at https://github.com/viktorZenkov/measuringAttraction/tree/master/Measuring. In addition, we provide 2 Imaris files that contain the movies from one small clustered/unclustered experiment and one large clustered experiment, featuring the “Spots” tracks of all T cells and the parasite in the imaging volumes (https://doi.org/10.5281/zenodo.5715658).

### 2.2 Metrics

To measure T cell bias (attraction or repulsion) towards the parasite, we define the following 4 metrics (Figure S3).

1. Angle metric (metric 1). For every movement of a cell, we calculate the angle between the cell’s movement angle and the angle to the parasite (Figure S3A). An acute angle corresponds to the T cell “getting closer”, and an obtuse angle corresponds to the T cell “getting farther”. For an unbiased cell making *n* movements, the choices of closer/farther for the movements are given by a binomial distribution with *p* = 0.5. This metric has been used extensively in ecology^19^.
2. Distance metric (metric 2). For every movement of a cell, we calculate *D*0, the change in the distance between the cell’s current position and the parasite, and *D*1, the distance between the cell’s next position and the parasite, with the change in distance designated by *r* = *D*0− *D*1 (Figure S3B). A negative change in distance corresponds to “getting closer” and positive corresponds to “getting farther. For an unbiased cell making *n* movements, the choices of closer/farther for the movements are given by a Poisson Binomial distribution with *p* = 0.5 − *r/*4*x*, where *r* is the cell’s movement length and *x* = *D*0 is the initial distance between the cell and the parasite, or *p* = 0 when *r* > *x* (Figure S4)^24^. This choice of null distribution is a critical improvement that we made which is necessary for this test to detect attraction at its fullest strength, which is explained further in the Supplemental information.
3. Angle distribution metric (metric 3). For every movement of a cell, we calculate the angle between the cell’s movement vector and the vector to the parasite and compare all angles with the von Mises-Fisher (vMF) distribution (Figure S3C and eqn. (1))^25,26^. By fitting the vMF distribution to the angle data, we calculate a concentration parameter *κ* which indicates the strength of a cell’s attraction towards the infection site, with *κ* > 0 indicating attraction and *κ* < 0 indicating repulsion. A 2D version of a similar, von Mises distribution-based metric, has been used in ecology^20^.
4. Average angle metric (metric 4). For every movement of a cell, we calculate the angle between the cell’s movement vector and the vector to the parasite^27^. We then use a Student’s t test to compare the mean of the angles to the expected average angle of 90 degrees with no attraction/repulsion^16,27^.

More information on the statistical tests using these metrics and the choices of null distributions, including thorough mathematical overviews demonstrating the validity of these tests, is provided in the Supplemental information.

### 2.3 von Mises-Fisher distribution

To quantify the degree of bias (or absence thereof) we introduce the von Mises-Fisher (vMF) distribution of angles towards the parasite^28^. The vMF distribution describes a probability distribution on an *n*-dimensional (we use *n* = 3) sphere given a direction vector and a concentration parameter *κ*. Sampling from the distribution gives a vector chosen pseudorandomly with a bias toward the given direction whose strength depends on *κ*. When *κ* → 0, the vMF distribution approaches a uniform distribution; *κ* > 0 indicates positive bias (attraction); and *κ* < 0 indicates negative bias (repulsion, Figure S5). We reduce the vMF distribution from a vector to a single angle between the output vector and the given vector - this angle *ϕ* corresponds to our angle metric. The probability density function of the angles of the vMF distribution with respect to the point of attraction is

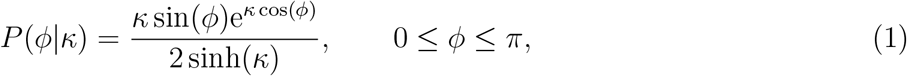

where *ϕ* is the angle between the vector to the attraction point and the cell’s movement vector. More intuitive parameters such as the fraction of acute angles *f*_*a*_ or the average angle towards infection 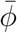 can be calculated as well (see eqn. (S4) in Supplemental information). Multiple biases, such as towards previous movement vectors or towards the infection site, can be naturally incorporated into a model using multiple vMF distributions (see Supplemental information).

To estimate the concentration parameter *κ*_*a*_ of a set of movement angles, we used the maximum likelihood approach (eqn. (S5)); more specifically, we used the `Maximize` function in Mathematica to estimate *κ*_*a*_. To evaluate if the estimated concentration parameter is statistically different from 0, we used a log-likelihood test by comparing the negative log-likelihood of the best fit and the negative log-likelihood with *κ*_*a*_ → 0. Details of how the vMF distribution can be used to simulate random walks with biases, a minor correction to the test similar to the improvement we made to the distance metric, and an analysis illustrating that the vMF distribution-based metric is the most powerful out of our tested metrics to detect attraction, are given in the Supplemental information (e.g., see Figures S6 and S7).

### 2.4 Simulations

To understand various aspects of T cell movement with respect to a parasite, we performed stochastic simulations using Mathematica 12.0. In our simulations, cell movements are characterized by a distance traveled per time step and a direction of cell movement (with respect to the previous movement and/or to the parasite). Movement lengths are chosen from a Generalized Pareto (Pareto type IV) distribution, and the direction of the movement is chosen from a vMF distribution with a given concentration parameter *κ*. Additionally, we developed a new method to simulate attracted cell movement using a variation of the Ornstein-Uhlenbeck process, one of the main methods used in biophysics to simulate correlated random walks^29^. Details of our simulation methods are provided in the Supplemental information.

## 3 Results

### 3.1 Only a small proportion of activated liver-localized CD8 T cells display attraction towards malaria liver stages

To determine if liver-localized CD8 T cells display attraction towards malaria liver stages, we performed novel experiments in which we tracked positions of malaria-specific CD8 T cells, CD8 T cells with irrelevant specificity, and malaria liver stages over time in murine livers using intravital microscopy (see Materials and Methods for more details). We attempted to design our experiments to detect s “first contact” event in which the first T cell locates the infection site, and such an event did occur in one experiment (described in section 3.3). In all other experiments, by the starting time of imaging, a small (1-2) or large number (5-7) of T cells had already formed a cluster around the liver stage (Figure S9). In general, imaging experiments performed further after the sporozoite infection resulted in larger clusters (results not shown). Notably, T cells displayed different movement characteristics depending on whether there were no/few or many cells in the cluster (Figure 1i-ii). Specifically, in unclustered/small clustered data, cell speeds were 2.42 ± 2.62 and 2.63 ± 3.03 *μ*m*/*min for OT1 and P14 cells, respectively (mean ± standard deviation, Figure S10). In the large clustered data, both cell types were significantly slower, with speeds of OT1 and P14 cells being 1.55 ± 1.66 and 1.59 ± 1.30 *μ*m*/*min, respectively (Mann-Whitney test, *p* < 0.001). In the presence of large clusters, T cells were likely to have a larger arrest coefficient (fraction of cell movements with a speed below 1 *μ*m/min): 0.29 and 0.27 for OT1 and P14 cells, respectively, in unclustered/small cluster data vs. 0.47 for both cell types in large cluster data. Additional analysis based on the meandering index (the distance between the first and last recorded positions divided by the total length of the path) and turning angles for T cells showed that all cells tend to turn (Figure S11) suggesting an active search for an infection.

**Figure 1:**
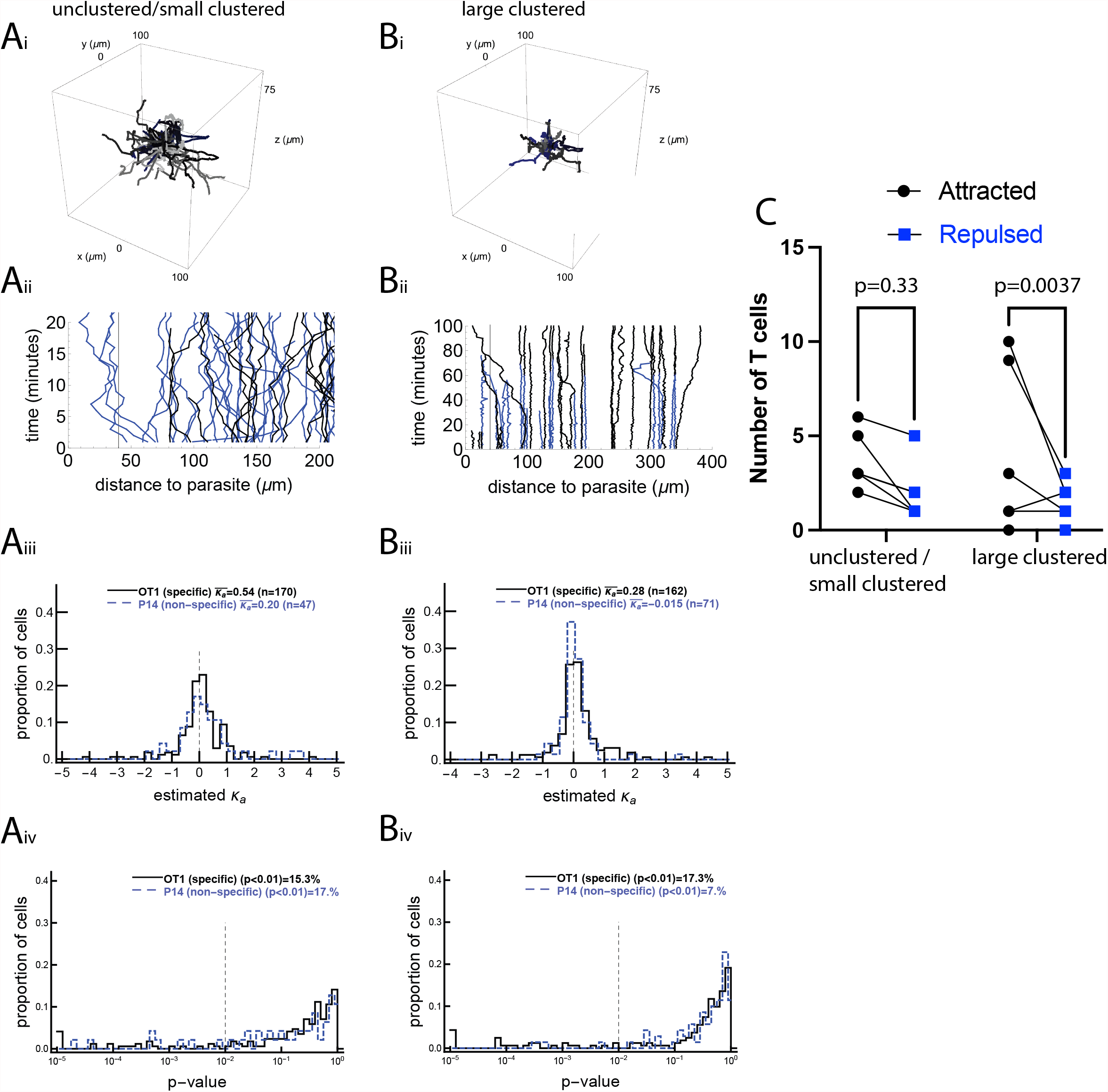
A minority of activated CD8 T cells display movement bias to the malaria liver stage. We performed four sets of experiments in which the movement of CD8 T cells with respect to the location of the malaria liver stage was recorded using intravital microscopy (see Materials and Methods), and performed tests on this movement data. In the unclustered/small clustered dataset, the movement of malaria-specific (OT1) CD8 T cells and T cells of irrelevant specificity (P14) was recorded around a single liver stage of Pb when no or few T cells were near the parasite (dataset #1, panels A, 4 movies in total). In the large clustered dataset, the movement of malaria-specific (OT1) CD8 T cells and T cells of irrelevant specificity (P14) was recorded around a single liver stage of Pb when several T cells were already near the parasite (dataset #2, panels B, 3 movies in total). Panels i show the T cell tracks from one of each set of experiments (the second infection site of the unclustered/small clustered data and the first infection site of the large clustered data) and panels ii show the distance from each T cell to the liver stage over time for the same infection sites. For each T cell we calculated the bias of T cell movement towards the parasite by estimating the concentration parameter *κ*_*a*_ from the vMF distribution (see eqn. (1) and Figure S3) using the maximum likelihood method. Panels iii show the distribution of estimated *κ*_*a*_s and panels iv show p-values from the likelihood ratio test for the vMF distribution with 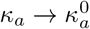, where 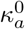 is the threshold value from the null distribution (see text for more detail of how 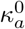 is calculated). The average concentration for all cells is shown on individual panels in iii, and the percent of T cells with a statistically significant bias to the parasite (with *p* ≤ 0.01 from our log-likelihood test) is shown in panels iv. Biased cells include both attracted (*κ*_*a*_ > 0) and repulsed (*κ*_*a*_ < 0) cells. In the unclustered/small clustered dataset, we detected 16 OT1 and 5 P14 cells as attracted and 10 OT1 and 3 P14 as repulsed. In the large clustered dataset, we detected 22 OT1 and 2 P14 cells as attracted and 6 OT1 and 3 P14 as repulsed. The number of OT1 cells detected as attracted was significant for the large clustered dataset and not significant for the unclustered/small clustered dataset (panel C). Results of the analyses for two other datasets are given in the Supplemental information (Figure S8).

To understand how T cells search for the infection site, we first pooled all track data from a given dataset into one set and determined if T cells display attraction towards the infection by fitting a vMF distribution to the data and estimating the concentration parameter *κ*_*a*_ (see Materials and Methods for more detail). We found that T cells displayed a statistically significant but weak attraction towards the parasite (*κ*_*a*_ = 0.077 (*p* = 3.8 ×10^−4^) and *κ*_*a*_ = 0.085 (LRT, *p* = 3.9 ×10^−5^) for the unclustered/small clustered and large clustered datasets, respectively, Figure 1); the concentration parameter *κ*_*a*_ = 0.077 corresponds to only 52% of movements being towards the parasite. There was stronger attraction of *Plasmodium*-specific CD8 T cells towards the infection site detected in the other two datasets with infections (*κ*_*a*_ = 0.19 (*p* = 0.003) and *κ*_*a*_ = 0.51 (*p* = 0.001), in Figures S8 and S12).

To investigate if weak attraction towards the parasite comes from all cells exhibiting weak attraction or a few cells exhibiting strong attraction, we calculated *κ*_*a*_ for individual cells. There was a broad distribution of *κ*_*a*_ for individual cells, but most of these values were not statistically different from a random value, with the average of these values indicating attraction (Figure 1iii-iv and Figure S8iii-iv). It should be noted that because the vMF distribution-based metric is inherently slightly biased for T cells moving with a correlated random walk, for every T cell track we calculated the null hypothesis concentration parameter 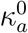 that would be expected given the cell’s initial position (with respect to the parasite and imaging volume), turning angle distribution, and movement lengths, and statistically compared the true *κ*_*a*_ to the null hypothesis value 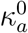 (results not shown). In most cases, this calculated null hypothesis 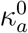 was not very different from zero. Notably, only OT1 cells (specific for malaria liver stages) in the large clustered data displayed statistically significant attraction towards the infection site (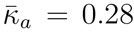; *p* = 0.0065, signed rank test), while P14 cells in the large clustered data and both OT1 and P14 cells in the unclustered/small clustered data did not display significant attraction (*p* > 0.05, signed rank test). Similarly, only malaria-specific T cells (PyTCR) displayed attraction (or weak attraction) for the two other datasets (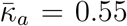; *p* = 0.0024 in the co-clustered dataset and 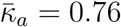; *p* = 0.059 for the Paris dataset), while non-specific cells (OT1 in the co-clustered dataset) were not attracted to the infection site (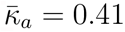; *p* = 0.13, Figure S8).

We found that about 15% of both malaria-specific and non-specific T cells displayed bias to the parasite (*p* < 0.01, Figure 1iv). Interestingly, while the fraction of *Plasmodium*-specific T cells (OT1) displaying bias toward the infection was similar between the unclustered/small clustered and large clustered datasets (15% vs. 17%, respectively), there were more P14 T cells (specific to LCMV) that displayed bias to the parasite in the unclustered/small clustered data than in the large clustered data (17% versus 7%, Figure 1iv), suggesting that some “biased” T cells in the unclustered/small clustered dataset may be an artifact of the statistical analysis. Indeed, we found similar fractions of T cells detected as attracted to or repulsed from the parasite in all cases except for the OT1 cells in the large clustered data (22/28, binomial test *p* = 0.004), suggesting that only during larger clustering is there a strong bias in (a minority of) malaria-specific CD8 T cells towards the infection site (Figure 1C).

### 3.2 Detecting bias to the infection is correlated with higher cell speed and persistence of movement

Our analyses suggest that the majority of CD8 T cells searching for malaria liver stages perform such a search without displaying a detectable bias towards the infection site. However, some malaria-specific CD8 T cells do display a detectable strong bias towards the infection site when there are already some T cells near the parasite. We next sought to determine which T cell characteristics may be correlated with bias towards the infection. To increase the power of the analysis, we pooled the data for malaria-specific CD8 T cells and CD8 T cells of irrelevant specificity. Previously it was suggested that the distance between an infection site and the T cell may determine the strength of attraction^16^. However, we found that the detected degree of attraction did not correlate with the starting distance between T cells and the infection (Figure 2A and Figure S13A). The distance to the closest T cell or the overall time per track also did not correlate with attraction (Figure 2D&E and Figure S13D&E), while the average movement per time step, cell velocity, and walk persistence (determined by the concentration parameter *κ*_*t*_) strongly correlated with the degree of attraction (Figures 2 and S13), suggesting that rapidly moving T cells are more likely to display bias towards the infection site. We also found that for the unclustered/small clustered data, velocity and small turning angles (the latter of which are represented by a large concentration parameter *κ*_*t*_) are correlated with cells displaying bias toward the infection site (Figures S14 and S15). This may be a statistical artifact because cells moving in a straighter path may randomly have their trajectory aim toward or away from the parasite, which would cause the cells to be detected incorrectly as attracted if the trajectory is toward the parasite and as repulsed if the trajectory is away from the parasite. Therefore faster and more persistent cells may be naturally inclined to be detected as biased toward the cell, but only as a result of chance.

**Figure 2:**
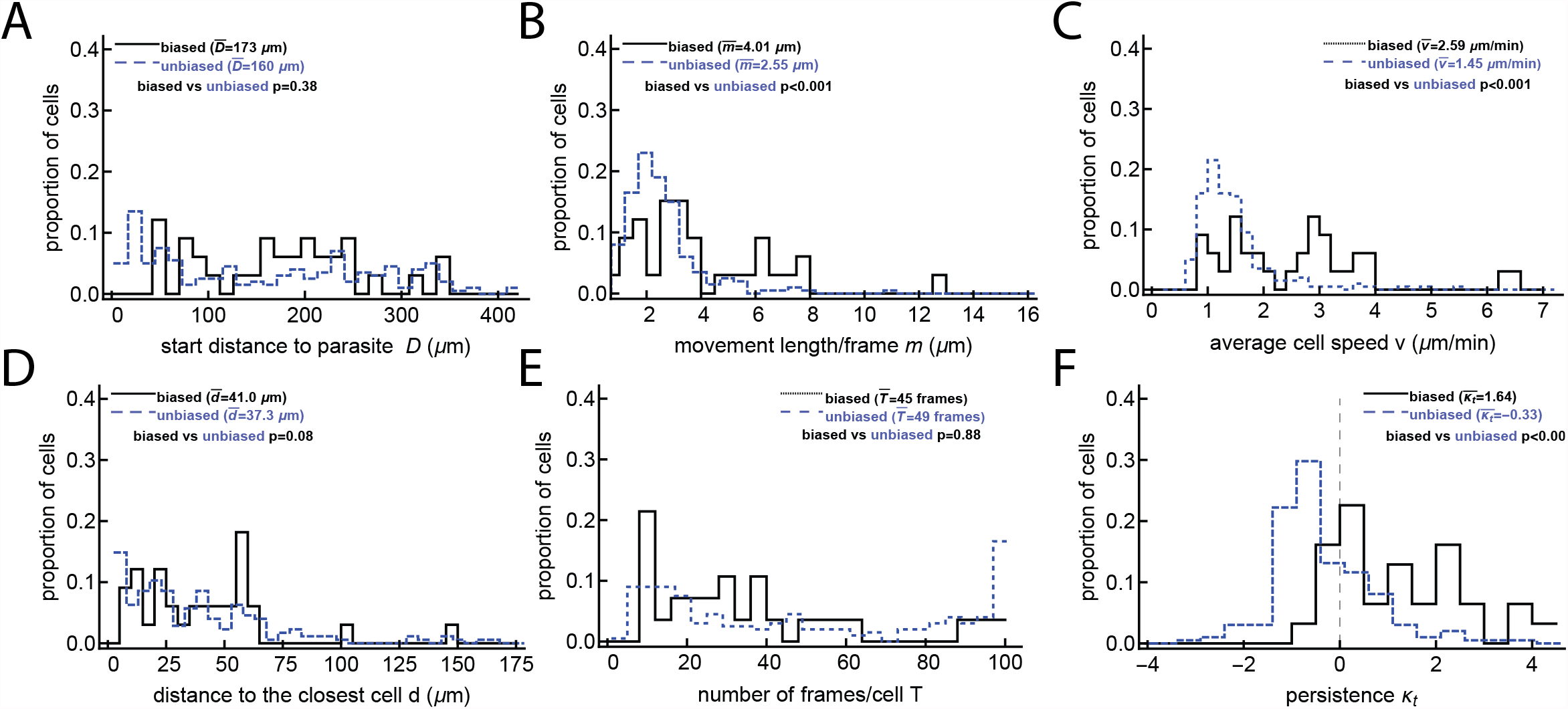
T cells detected as biased are correlated with faster T cells and T cells with smaller turning angles. For the large clustered dataset, in which some T cells have already located the parasite (shown previously in Figure 1B), we calculated multiple characteristics for cells that display no bias to the parasite’s location (“unbiased” cells) and cells that do display bias to the parasite’s position (“biased” cells, i.e. attracted or repulsed). These characteristics include: starting distances *D* (panel A), movement lengths per frame *m* (panel B), average speed per T cell *v* (panel C), distance to the closest T cell *d* (panel D), the number of recorded positions *T* (panel E), and the estimated concentration of the vMF for the turning angles *κ*_*t*_ (panel F). Comparisons were done using the Mann-Whitney test and the p-values for the comparisons are shown on individual panels. The unbiased results are offset slightly to not directly overlap with the biased results for ease of viewing. Analyses when the data were divided into attracted, repulsed, or unbiased T cells are shown in Figure S13, and analysis of the data with no/small clusters is shown in Figure S15. The majority of biased T cells in this dataset display attraction towards the infection (24/33, binomial test *p* = 0.014).

### 3.3 The amount of detected attraction does not noticeably change immediately upon formation of a cluster

Our results show a greater proportion of attraction toward parasites in data with large CD8 T cell clusters. This potentially suggests that the first cells to reach the parasite may do so randomly, and then other cells begin to show bias after the environment around the parasite has been changed by the first scout cells. In one of our experiments we did observe such a “first contact” event where, at the start of imaging, no T cells were near the parasite, but then 1 cell out of 61 in the imaging volume reached the parasite, and then 2 more cells entered the cluster and 1 cell left (Figure 3 and Figure S9Aii). We then analyzed whether those T cells, which were not in the cluster after the first scout T cell located the parasite, displayed bias towards the infection. We pooled the angles of T cell tracks after different times in the experiment and calculated the overall attraction towards the infection, characterized by the concentration parameter *κ*_*a*_ (Figure 3A) and the corresponding p value from the LRT (Figure 3B). Interestingly, the estimated *κ*_*a*_s for angles after an hour after the first scout T cell found the infection site were still not significantly different (Figure 3). A statistically significant repulsion (*κ*_*a*_ < 0) after later time points is most likely the result of noise because of very limited data after late time points. Our analysis suggests that if the first T cell that located the parasite changes the environment, such a change takes longer than ∼ 50 min to be detected by other moving T cells.

**Figure 3:**
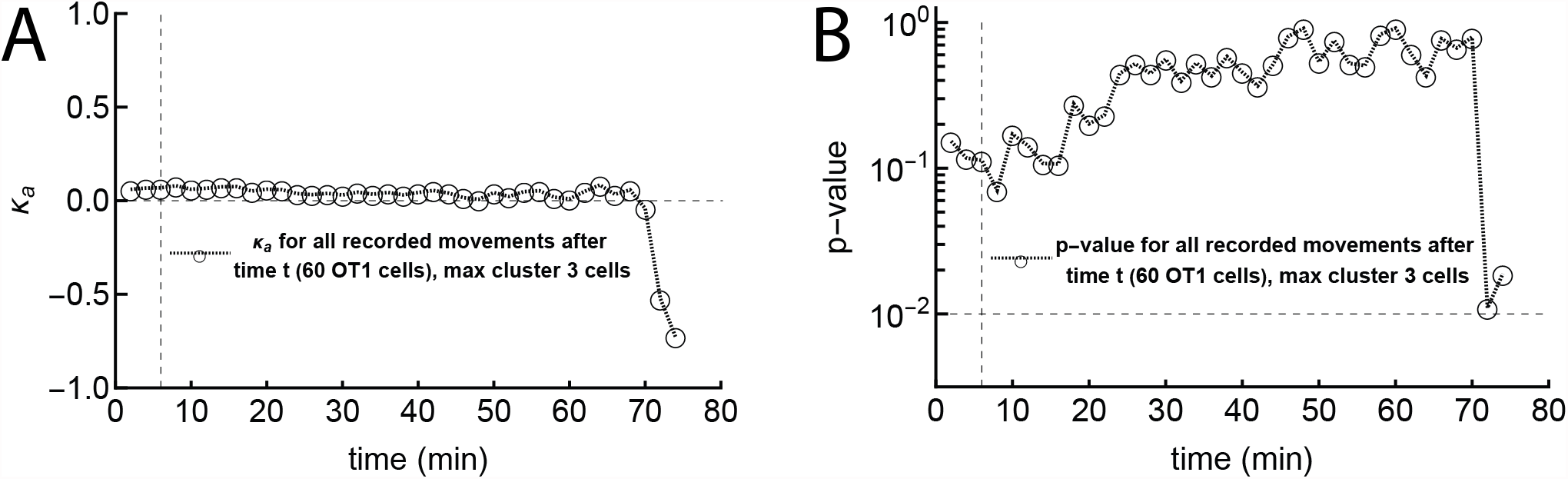
The first scout T cell locating the infection site does not impact the movement bias of other T cells towards the infection within an hour. In this “first contact” experiment (Figure S9Bii), no T cells were close to the parasite at the start of imaging, and the first cell found the parasite in six minutes (marked with a vertical dashed line in both panels). After this cell, two other cells reached the parasite within a few minutes, but no more cells (out of a total of 60 cells in the video) reached the parasite within an hour (and in fact, one of the three left the parasite). We calculated the concentration parameter *κ*_*a*_ for all T cell tracks combined after different times (panel A) and the corresponding p value from the likelihood ratio test (LRT) indicating deviation of *κ*_*a*_ from 0 (panel B). The horizontal line in panel B denotes p value of 0.01.

### 3.4 Detecting weak attraction is difficult with current experimental setups

In our analyses so far we found that, with some exceptions, the vast majority of liver-localized CD8 T cells search for the malaria liver stages randomly, with little evidence of bias towards the infection site. Yet, we know that with sufficient numbers of liver-localized CD8 T cells, all liver stages will be eliminated within 48 hours^7,30^. We reasoned that while our liver imaging experiments were sufficiently long (∼ 1.5 − 2 hours) to detect attraction in some cases, it may be possible that they were still too short to detect any weak attraction displayed by individual T cells. Therefore, we performed several sets of simulations to determine the length of experiments that would be required to detect a given degree of T cell attraction towards the infection site (Figure 4).

**Figure 4:**
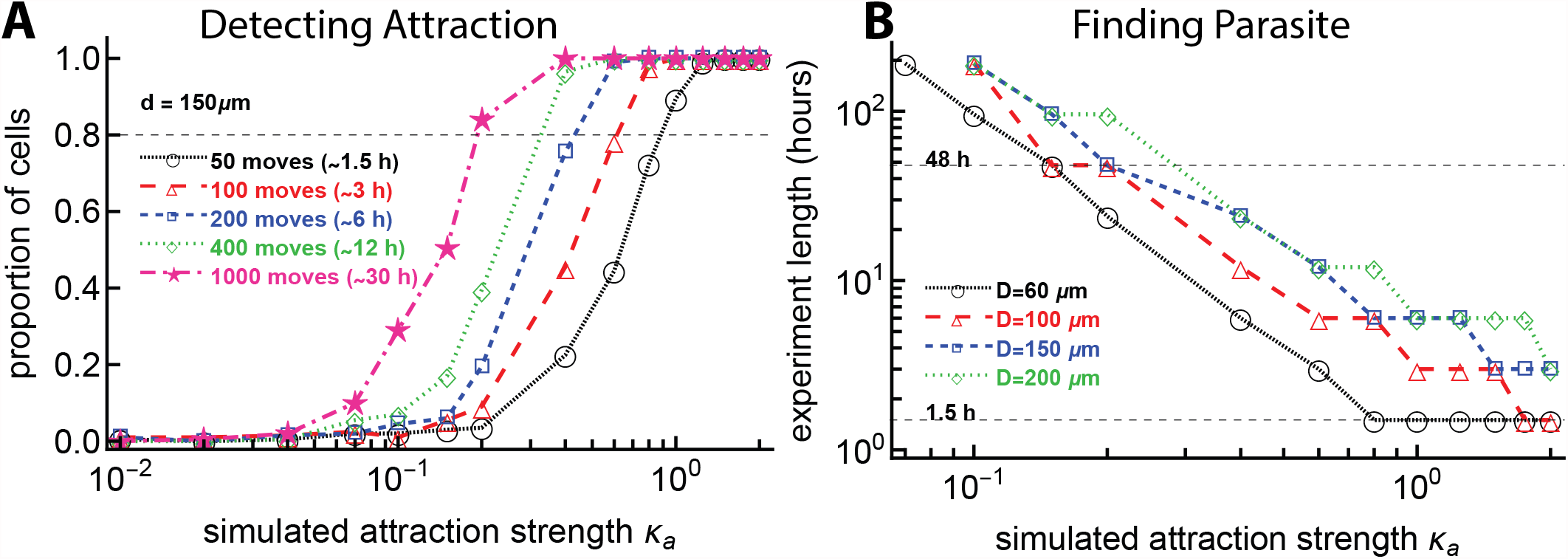
To detect weak T cell attraction to the parasite or to observe a T cell finding the parasite requires a prohibitively long imaging duration. In panel A we simulated T cell movement with some degree of attraction to the parasite defined by the concentration *κ*_*a*_ and starting at a distance *D* = 150 *μ*m. For 1000 simulated T cells, we varied the number of T cell movements (which changes the duration of the imaging movie). For every track we then determined the probability that the movement track would exhibit attraction (we estimated *κ*_*a*_ from the simulated tracks using maximum likelihood and used the likelihood ratio test to calculate if the estimated value was significantly different from zero, i.e. from random). In panel B we performed simulations with 1000 cells for values of *κ*_*a*_ between 0 and 2, with starting distances of 60, 100, 150, and 200 *μ*m from the parasite and T cell movement lengths chosen from a generalized Pareto distribution (eqn. (S8)) with pseudorandomly chosen turning angles. We calculated the time it took for the T cell to reach the parasite (reach a distance of 40 *μ*m from the parasite). A typical imaging duration (∼ 1.5 h) and the time it takes for liver stages to develop (48 h) are indicated by dashed horizontal lines in panel B.

We simulated movement of 1000 cells for combinations of varying strengths of attraction towards the infection site (*κ*_*a*_) and numbers of cell movements (choosing numbers of movements with an 2 min timestep, imitating the real data), assuming that cells start their search at 150 *μ*m from the parasite. For each combination of parameters, we determined the power to detect attraction as the proportion of cells which are detected as attracted based on the angle distribution (vMF distribution-based) metric. For our typical experiment with a length of 1.5 hours (about 50 movements of T cells), the attraction strength *κ*_*a*_ must be more than 1.0 for the bias of T cell movement towards infection to always be detected (Figure 4A). Furthermore, to detect weaker attraction (*κ*_*a*_ = 0.2), 30 hours of imaging experiments (or 1000 movements per cell) would be needed, which is not currently possible due to ethical and experimental constraints. The need for 30 hours’ worth of data suggests that our failure to detect weak attraction of T cells towards the infection site may be due to limited (but the best currently possible) data.

We wondered how the time to find the parasite may depend on the starting distance from the T cell to the infection site and the degree of the T cell attraction towards the infection site. To answer this, we performed another set of simulations (with 1000 cells for each combination of parameters) by varying the starting distance between the T cells and the parasite and the degree of T cell attraction to the parasite (*κ*_*a*_). We then calculated the time when a cell found the parasite (reaches within 40 *μ*m from the parasite) with 80% probability. A high degree of attraction (*κ*_*a*_ ∼ 1) and short distance is required for T cells to find the parasite within 1.5 hours, and to reach the parasite in 48 hours still requires moderate attraction (*κ*_*a*_ = 0.3, Figure 4B). For a T cell with *κ*_*a*_ = 1.0 and a starting distance of 150 *μ*m, the experiment must last for 6 hours for the T cell to reach the parasite; currently we are not able to perform experiments with live mice for this length of time. These findings suggest that there may be something special about these parasites that had T cell clusters form around them early after the infection. For example, such parasites may be entering the liver where there are already some T cells nearby, perhaps indicating that *Plasmodium* infection of the liver and T cell localization in the liver do not occur randomly. Furthermore, analyses also suggest that some level of weak attraction is needed to explain T cells finding the parasite in 48 hours after infection.

### 3.5 Only a small fraction of T cells display bias towards the infection site

In our clustered data we found that only about 20% of T cells display statistically significant bias in movement towards the infection site (Figure 1iv). An alternative interpretation of this result is that perhaps all T cells in these experiments exhibit a small bias towards the infection site, but only a small fraction of the cells display detectable significant bias. To test this interpretation we performed additional simulations. Specifically, we simulated movement of 500 cells by varying the strength of attraction towards the infection site (*κ*_*a*_) for all cells and the number of cell movements (assuming that every movement was recorded with 2 min timesteps), and assuming that cells start their search at 150 *μ*m from the parasite. For each combination of parameters, we fit a vMF distribution to each of 500 trajectories, generating a distribution of concentrations *κ*_*a*_. One interesting observation from these simulations is that as the bias towards simulated infection sites increases, the mean of the estimated concentration parameters increases as well, leading to a large difference between experimentally observed and simulated distributions (Figure 5A-B, Kolmogorov-Smirnov test *p* < 0.001). The simulated distributions typically had smaller tails than the experimental distribution, implying less variation in the estimated concentrations *κ*_*a*_ (Figure 5A-B). This may partly be due to the simulated cells not having persistence in their movement, while real cells do appear to move with persistence that may lead a cell to move randomly in a continuous direction toward or away from the parasite. The differing sizes of the tails suggest that our data are not consistent with the idea that all cells exhibit some level of bias towards the infection site. Furthermore, power analyses suggest that to see at least 20% of cells exhibit statistically significant bias towards the infection, all cells must have a relatively substantial attraction (at least *κ*_*a*_ ≈ 0.5, Figure 5D). That consistently high attraction would result in a high average attraction not observed in actual data. Thus, our results are more consistent with the explanation that only a small fraction of T cells exhibit bias towards the parasite when there is already a cluster of T cells.

**Figure 5:**
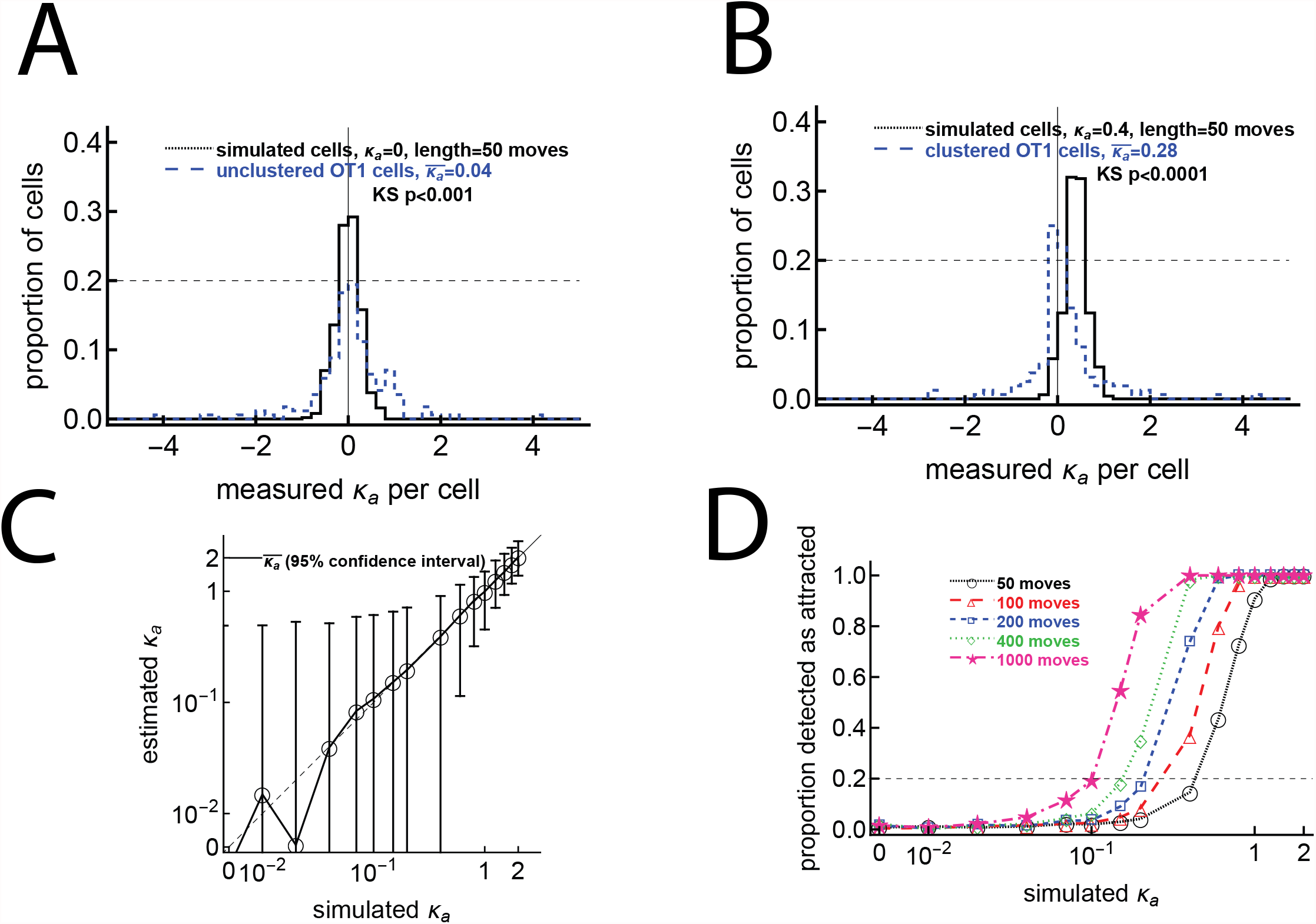
The hypothesis that all cells have weak attraction is not supported by the data. We simulated CD8 T cells searching for a malaria liver stage assuming variable levels of T cell attraction to the parasite, defined by the concentration parameter *κ*_*a*_ of the vMF distribution and with different number of movements (from 50 to 1000 movements/cell). For each cell we calculated the observed attraction towards the infection site using the vMF distribution (eqn. (1)). In panels A and B we overlay distributions of detected *κ*_*a*_ from simulated and actual datasets. Panel A shows data for unclustered cells and simulated cells with no inherent attraction (*κ*_*a*_ = 0.0001), while panel B shows data for clustered cells and simulated cells with a minor attraction (*κ*_*a*_ = 0.1). P values using the Kolmogorov-Smirnov (KS) test to compare the experimental and theoretical distributions are shown on the panels. Panel C shows the mean estimated *κ*_*a*_ from 500 simulated cells for the simulations with 50 moves and each original *κ*_*a*_; the dashed line denotes the curve *y* = *x*; and error bars denote 95% confidence intervals on the estimated *κ*_*a*_. Panel D shows the proportion of cells detected as attracted for each combination of simulated *κ*_*a*_ and number of movements; the horizontal dashed line denotes 20% of cells detected as attracted as we have observed in our experimental data. In simulations, all cells start 150 *μ*m from the parasite. To simulate a T cell walk we assumed that T cell movement lengths follow a generalized Pareto distribution (eqn. (S8)) with pseudorandomly chosen turning angles. Simulations were done for 500 cells for each value of *κ*_*a*_ and experiment length.

### 3.6 T cells only display exclusive attraction to the infection site when a T cell cluster has already formed around the parasite

In our analysis, we focused on detecting if T cells are attracted to the site of infection, and our tests did not compare *Plasmodium*-specific cells with control T cells (specific to an irrelevant antigen) because those results could be biased if both malaria-specific T cells and T cells with irrelevant specificity may be equally attracted to the infection site. However, our vMF distribution-based metric could be too sensitive, detecting attraction when it does not exist. Indeed, we showed that for T cells moving with a correlated random walk, some cells may be detected as attracted to the infection in the absence of actual attraction (Figure S6). We reasoned that if T cells are truly attracted to the infection site, them fewer cells (if any) should display attraction to areas distant from the actual parasite’s position.

For this analysis we chose “fake parasite” positions equally spaced in the 500 ×500 ×50*μ*m^3^ imaging box (a total of 216 positions) and determined the number of real T cells that were detected as attracted to (or repulsed from) each fake parasite location (tested using our vMF distribution-based metric). For the no parasite (uninfected mouse) and unclustered/small clustered (infected mice) datasets, the number of cells detected as attracted to (or repulsed from) any position was similar. This implies that the detection as attracted or repulsed is simply a statistical artifact. The former option seems unlikely, so we surmise that in these datasets, any attraction or repulsion detected to the real infection site is simply an artifact (Figure 6i-ii). This corroborates the conclusion that in the absence of large cluster formation, T cells were searching for the parasite randomly (or with a weak bias that was not detectable). In contrast, in the large clustered dataset, more cells are detected as attracted and fewer as repulsed for fake parasite positions around the real parasite (for which the distance between the real parasite and “fake” parasite is small), suggesting that there truly is some T cell attraction to the real parasite position when there are other T cells already present near the parasite (Figure 6iii).

**Figure 6:**
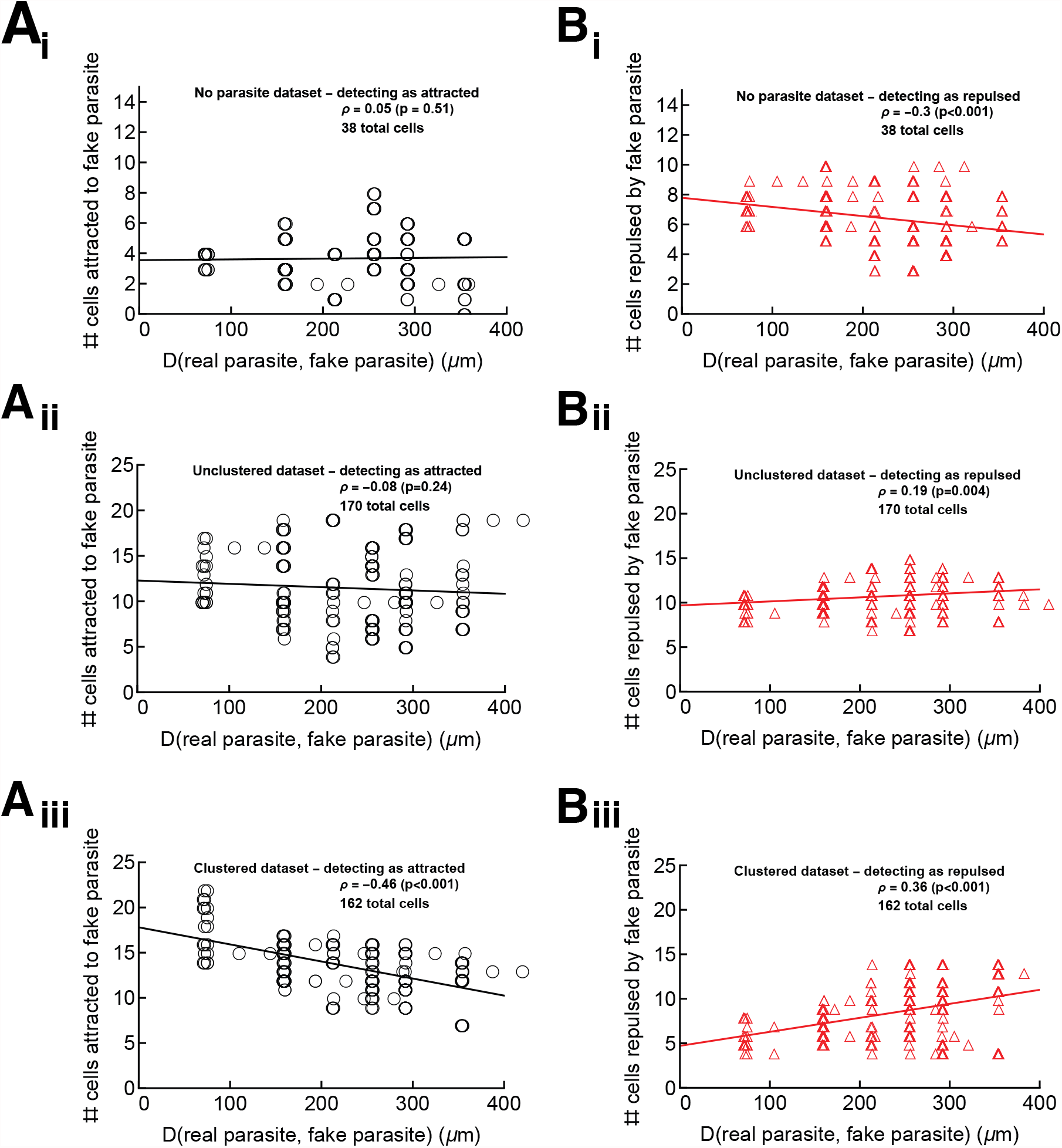
The parasite’s position is an attraction point for T cells only in the large clustered data. We performed analyses to test if cells in the “no parasite” data (panels i), in the unclustered/small clustered datasets (panels ii), and in the large clustered datasets (panels iii) display attraction to sites other than the infection site. We chose positions of a “fake” parasite every 10 *μ*m on the grid extending 250 *μ*m away from the real parasite. For the “no parasite” movie, which has no infection, the “true” parasite was assumed to be in the center of the imaging volume for the purposes of judging the distance of fake parasites from the true parasite. Then for every T cell we calculated the concentration parameter *κ*_*a*_ (using eqn. (1)) and determined if the estimated parameter is different from 0 using LRT (see Materials and Methods for more detail). In this way, we calculated the number of T cells attracted to (*κ*_*a*_ > 0, panels A) or repulsed from (*κ*_*a*_ < 0, panels B) the “fake” parasite and the distance *D* between each “fake” parasite position and the true parasite position (denoted on *x* axes in the panels). The changes in the numbers of cells detected as attracted or repulsed with distance *D* were tested using the Spearman rank correlation test (the rank correlation *ρ* and p values from the tests are indicated on individual panels). The lines from linear regression are shown for visual purposes only.

### 3.7 A lower number of T cells is required for sterilizing protection if there is biased T cell movement towards the infection site

If CD8 T cells search for the malaria liver stages nearly randomly, we wondered how the presence of weak attraction (that is not detectable in current experiments) could change the ability of T cells to find the parasite (i.e., reach within 40 *μ*m of the parasite). We performed 1000 simulations for each combination of varying T cell attraction to the parasite (the concentration *κ*_*a*_) and starting distance from the cell to the parasite, and counted the cells which reach the parasite (within 40 *μ*m) within a defined time period. Our results suggest that a small change in attraction strength *κ*_*a*_ can dramatically increase the chance of T cells finding the parasite, especially when the initial distance between the T cell and parasite is large (Figure 7A-C). Importantly, a relatively weak bias towards the infection (at least *κ*_*a*_ ≈0.2 − 0.3) is sufficient for T cells to locate the parasite within 48 hours after infection, even for a relatively large initial distance between the T cell and the parasite (Figure 7C).

**Figure 7:**
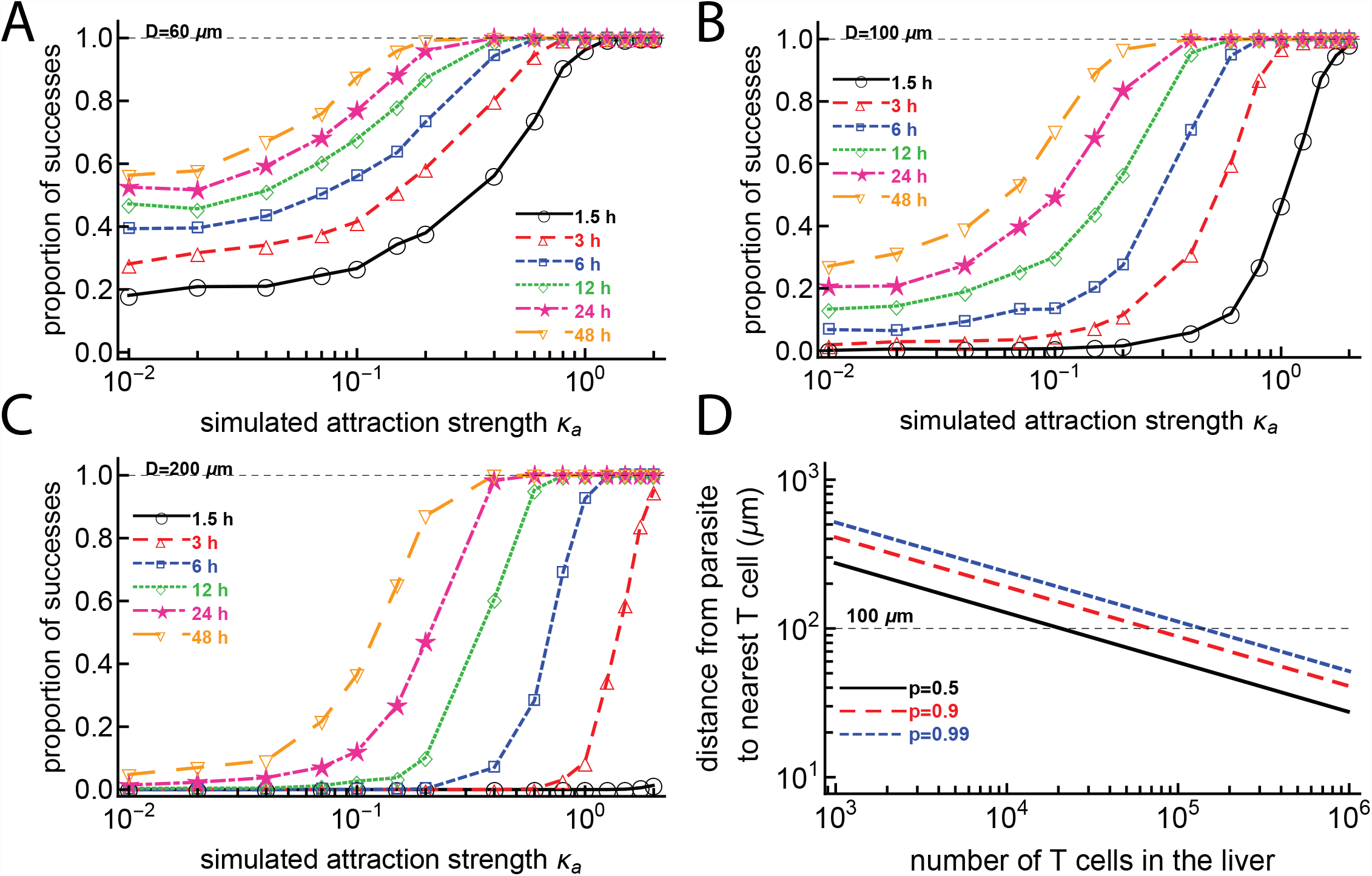
Small attraction dramatically increases the chances for a T cell to find the parasite within 48 hours. We simulated CD8 T cells searching for a malaria liver stage assuming variable levels of T cell attraction to the parasite, defined by the concentration parameter *κ*_*a*_ of the vMF distribution. Cells started their search for the infection 60 (A), 100 (B), or 200 (C) *μ*m from the parasite. We calculated the probability that the cells reach the parasite within 40*μ*m distance at various times after infection. To simulate a T cell walk we assumed that T cell movement lengths follow a generalized Pareto distribution (eqn. (S8)) with pseudorandomly chosen turning angles. We did simulations for 1000 cells for each value of *κ*_*a*_. In panel D we calculated the distance between a parasite and the nearest CD8 T cell assuming T cells are randomly distributed in the 1 mL = 1 cm^3^ volume of the liver, and showed different levels of certainty that the cell is within that distance (defined by the probability *p*). The distance from the parasite to the nearest T cell is calculated as 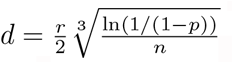, where 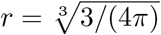 cm is the radius of a sphere with volume 1 cm^3^ (this approximates a mouse liver) and where *n* is the number of T cells in the liver. For *p* = 0.99 when there are *n* = 10^5^ T cells in the liver, the chance that one T cell is at the distance of *d* = 100 *μ*m from the parasite is 99%.

In our experiments, the initial distances between T cells and the parasite varied dramatically; the average was around 150-200 *μ*m (e.g., Figure 2). Given a mouse liver volume of about 1 mL^31^, we calculated that if there are 10^5^ randomly distributed T cells in the liver (a number found experimentally to provide sterilizing protection against malaria in mice^32,33^), the distance between a randomly positioned parasite and its nearest T cell is about 100 *μ*m, with 99% probability (Figure 7D). However, without biased movement towards the parasite, CD8 T cells located at 100 *μ*m from the parasite are unlikely to locate the infection even within 48 hours (Figure 7B), suggesting that some weak attraction may be guiding T cell search for the infection.

Our simulations so far focused on scenarios where a single T cell reaches the parasite. However, when multiple T cells search for the parasite, the probability of at least one T cell finding the infection could be higher. To investigate this, we performed 1000 simulations to allow several T cells to search for the parasite by varying the strength of T cell attraction to the parasite (concentration *κ*_*a*_), the number of searching T cells, and the length of search, and by fixing the starting distance of T cells from the parasite to 150 *μ*m. For each combination of parameters, we determined the proportion of the 1000 simulations for which at least one T cell reaches the parasite (within 40 *μ*m). Results suggest that for essentially all parasites to be found in 48 hours, we need around 10 T cells per parasite and a weak attraction defined by a concentration of at least *κ*_*a*_≈ 0.2 (Figure S16). These findings suggest that when multiple T cells search for an infection, weak attraction greatly improves the chances of T cells finding the parasite.

## 4 Discussion

It has been well established that activated and memory CD8 T cells are capable of providing sterilizing protection against infection of *Plasmodium* sporozoites in mice^7,9,30^. However, the ways that T cells locate and eliminate all *Plasmodium*-infected hepatocytes remain unclear. By generating unique data on the movement of liver-localized activated CD8 T cells within a few hours after infection with *Plasmodium* sporozoites, and by rigorous analyses of these data using a newly developed metric based on the von Mises-Fisher distribution, we found that most CD8 T cells search for the infection site randomly. Using stochastic simulations, we showed that randomly searching T cells have a high failure rate of finding parasites. A small bias towards the infection dramatically improves the chance of T cells finding the infection, and thus reduces the number of liver-localized T cells needed for sterilizing protection. Power analyses showed, however, that current experimental set-ups allowing for intravital imaging of livers of live mice for only a few hours do not generate sufficient data to detect attraction to the infection site by individual T cells.

For all analyzed datasets, we consistently found that a small fraction (about 15-20%) of T cells displayed strong movement bias to the infection site; such biased T cells included both T cells attracted to the parasite and T cells repulsed from the parasite. In the case where there were several T cells near the parasite (large clustered data), the number of T cells attracted to the infection site was significantly higher than that of repulsed cells. The distance between the T cell and the infection did not correlate with the strength of attraction; however, the speed at which T cells moved was a strong predictor of attraction, that is, more rapidly moving T cells displayed stronger bias. This observation highlights the need for a better understanding of the mechanisms that regulate the movement speed at which T cells survey peripheral tissues. In our case, however, we found that rapidly moving T cells having a higher bias towards the infection site may be an artifact of a correlated/persistent random walk.

The reasons why some T cells (e.g., OT1 cells in the large clustered data) display attraction towards the infection site while others (e.g., P14 cells in the large clustered data) search randomly are unclear. It is generally believed that movement patterns of T cells in tissues are regulated by chemokines, i.e., T cells with appropriate chemokine receptors follow chemokine gradients in the environment^34^. However, it is not yet understood if T cells indeed follow chemokine gradients (which may be shallow in many circumstances) or if chemokines simply regulate the cells’ velocities^16,34,35^. For example, CXCL21 chemokine and CCR7 receptors on B cells regulate the localization of B cells near B-T zones of lymph nodes^35^. However, while the CXCR3 receptor is important for CD8 T cell movement in the skin or brain^16,36^, whether T cells in these tissues follow the chemokines’ gradients has not been established. We recently found that the LFA-1 receptor on T cells is critical for T cell motility in the liver; however, even LFA-1-deficient T cells are capable of locating *Plasmodium* liver stages with a somewhat lower success rate^23^. Whether receptors regulate T cell attraction to the *Plasmodium* liver stages is unclear. It is possible, however, that such attraction is achieved by a combination of receptors, and the lack of any single receptor does not dramatically impact the ability of T cells to locate the infection. It was recently found that the migration of *Mycobacterium tuberculosis*-specific CD4 T cells from the lung vasculature into parenchyma is dependent on many receptors, and lacking a single receptor had only a moderate to minimal impact on the rate at which T cells migrate from the blood to the lung^37^.

The presented results have several limitations. Even though our intravital imaging experiments lasted for 2-3 hours, the amount of data collected was not sufficient to detect the weak attraction that individual T cells may exhibit when searching for the parasite. Experiments with a longer duration may be ethically difficult, and long-term imaging of surgically exposed livers may also generate artifacts due to tissue damage. The imaging frequency in our data was relatively low (1.5-2 min per stack of images) in order to allow for longer movies with lower laser exposure to the liver. While more frequent measurements could increase the number of data points, long movies may increase chances of tissue damage, require time-consuming processing steps, and result in a bias of imaged cells remaining in the imaging volume for a long time. Therefore we aimed for a balance between the length of imaging and the costs associated with longer movies.

Due to the limitations of intravital imaging, we could only record cell positions in a limited 3D volume of approximate 500 ×500 ×50 *μ*m^3^ centered near a parasite. If a cell exits that volume we no longer record its positions, so there may be bias toward recording more movements from T cells that act attracted to the parasite than cells that act repulsed from the parasite, since repulsed cells are more likely to leave the imaging volume. We performed simulations that approximately allow us to correct for such bias in our vMF distribution-based metric.

While it is not often mentioned, intravital imaging may induce local damage due to exposing tissues to lasers, especially in longer experiments. It is often difficult to evaluate the degree of liver damage any specific experiment involves. In our experiments we observed that in *Plasmodium* sporozoite-infected livers, CD8 T cells move relatively slowly, with speeds of 1.5-2.5 *μ*m*/*min. This was significantly slower than the 3.5 *μ*m*/*min estimated in one control movie without an infection (“no parasite” dataset), and lower than the average T cell speeds estimated in previous studies, which are in the range of 5-10 *μ*m*/*min^23,38^. One potential explanation of this difference may be due to the frequency of imaging, which in our experiments was rather low; evidence that imaging frequency has a direct influence on the inferred speeds of cells will be presented in a future work.

For some of our analyses, we pooled together the movements of different cells found in different mice to increase the analyses’ power. In general we found similar results (although at times statistically not significant) as when we analyzed the individual datasets separately. The vMF distribution, while a straightforward representation of bias in 3D movement, may not perfectly approximate the distribution of angles towards the infection, although it is close (Figure S12). We found that in cases when the vMF distribution does not fit the angle to infection data well, a mixture of vMF distributions can match the data very closely (Figure S12A-C). Such an approach also allows us to estimate the fraction of angles near 90^*°*^ that deviate from the single vMF distribution; what such angles represent biologically has yet to be determined. For example, this pattern may arise if cells attempt to approach a particular location and are unable to do so directly due to the absence of a direct path to the location; in this case the cells may thus meander.

Most of our simulations were concerned with the process of T cells finding the parasite; however, to protect from malaria, T cells must also kill the parasite. The time it takes for CD8 T cells to kill *Plasmodium* liver stages after reaching them has not been quantified, and therefore our estimates of the search time should be considered minimum estimates of the time it takes for T cells to find and kill the parasite.

One major limitation is that we did not take into account the fact that liver-localized CD8 T cells move in liver sinusoids^23,39^ and are thus constrained in their movement. The chance of randomly finding an infection may be influenced by the structure of the sinusoids^39^, and it remains to be determined how a biased search will find parasites in a constrained sinusoid structure instead of the open (3D) space in which all the metrics in this paper operate. An argument could be made that ignoring liver sinusoids makes the simulations not useful for approximating cell movement; however, as we choose simulation parameters to replicate real movement, we think our simulations, even though performed in open space, are an acceptable representation of cell movement in the liver. How CD8 T cells may search for the infection site in the constrained liver environment of sinusoids will be presented in a future work.

One of the purposes of this work was to test predictions of our recently published density-dependent recruitment model of CD8 T cell cluster formation around *Plasmodium* liver stages, which states that the formation of clusters is driven by a positive feedback loop where larger clusters recruit more T cells^10,11^. While the data are supportive of a random search of T cells for the parasite, it is still unclear why some parasites already have T cells nearby at the start of imaging while other parasites have no T cells nearby or there is no accumulation of T cells near other parasites in ∼1 − 2 hours of imaging. Even in situations in which the first T cell discovers the parasite, we found no evidence that T cells in surrounding areas become more attracted to the parasite within an hour. The data thus do not yet allow us to discriminate between models with density-dependent recruitment and with variability in attraction to parasites^11^.

There are several ways the work outlined in this paper can be extended. We did not investigate if any specific chemokine receptors (e.g., CXCR3, CCR5, or CX3CXR1) impact the success rate of the T cell search for the infection site. This is one focus of our current research. Including the structure of liver sinusoids in simulations of T cell searches for infection would likely provide more realistic estimates of the time it takes for T cells to find the parasite. To measure T cell attraction to the infection in this paper, we used the vector from the T cell to the parasite as the optimal route to reach the infection site. However, due to physical constraints from the liver sinusoids, the optimal route to the infection may not be the direct vector. By imaging the parasite, T cells, and liver sinusoids, it may be possible to quantify an optimal route, traveling through the sinusoidal structure, and to determine if the cells move with attraction along the structure. Work on this project is underway.

Future work may need to investigate how quickly T cells form clusters around the parasites and if the degree of attraction we estimate in this work is sufficient to explain the formation of relatively large clusters (e.g., 5-10 CD8 T cells) near individual parasites^10,11^. Similarly to how we used statistics to predict the distribution of angles to the parasite using the vMF distribution, we also attempted to find an analytical description of a distribution of changes of distances to the parasite with T cell movement. However, that distribution is more complicated mathematically, and we left it for future work. Our analysis propounds that the speed at which T cells search for an infection may be correlated with the ability of T cells to locate the infection site. However, there appears to be a trade-off between the speed T cells are moving in tissues and the ability of such T cells to process environmental signals to detect the infection - too rapid cells may miss many of the infected cells in their haste^40^. Whether there is an optimal movement pattern of T cells to locate all infections in a small enough time to cure infection is also work for a future project^41^. Despite potential limitations, our work provides novel data from innovative imaging experiments and rigorous mathematical analyses that begin to elucidate how CD8 T cells search for infection in the complex tissue of the liver.

## Data sources

T cell movement/track data generated in this paper (unclustered/small clustered, large clustered, and no parasite datasets) and published previously (Paris and co-clustered datasets) are provided in xlsx and csv format at (https://github.com/viktorZenkov/measuringAttraction/blob/master/Data/AllPositionData.xlsx and https://github.com/viktorZenkov/measuringAttraction/blob/master/Data/AllPositionData.csv). Also, we provide two Imaris files that contain original imaging data and objects (Spots) for unclustered/small clustered and large clustered datasets (https://doi.org/10.5281/zenodo.5715658).

## Code sources

Analyses have been primarily performed in Mathematica (ver 12) and codes used to generate most of the figures in the paper are provided on GitHub: https://github.com/viktorZenkov/measuringAttraction/tree/master/Measuring.

## Ethics statement

All animal procedures were approved by the Animal Experimentation Ethics Committee of the Australian National University (Protocol numbers: A2016/17; 2019/36). All research involving animals was conducted in accordance with the National Health and Medical Research Council’s Australian Code for the Care and Use of Animals for Scientific Purposes and the Australian Capital Territory Animal Welfare Act 1992.

## Author contributions

VSZ, JOC, IAC, and VVG developed ideas for the study. JOC performed intravital imaging experiments and segmentation of the imaging data with Imaris (under supervision of IAC). VSZ performed analyses of the data and simulations of cell movements (under supervision of VVG). VSZ wrote the first draft of the paper, with modifications later by VVG and VSZ with some contributions from JOC. VSZ is the primary author of the paper.

## Acknowledgments

We would like to thank Harshana Rajakaruna for discussion of various aspects of this work, and all members of GanusovLab for feedback on earlier versions of the paper. This work was supported by NIH grant (R01 GM118553) to VVG.

## Abbreviations

vMF: von Mises-Fisher
GFP: green fluorescent protein
TCR: T cell receptor
OVA: ovalbumin
LCMV: lympocytic choriomeningitis virus
LRT: likelihood ratio test
OU: Ornstein-Uhlenbeck.

## Supplemental Information

### Additional experimental details

Mice were prepared for two-photon microscopy as described in previous work^23^. In summary, mice were anesthetized with a mix of Ketamine (100 mg/kg) and Xylazine (10 mg/kg). Throughout the surgery and imaging pro cedure the mouse temperature was maintained at 37°C using a heating mat attached to a feedback probe inserted in the mouse rectum. A lateral incision was made over the left lobe of the liver and any exposed vessels cauterized by applying light pressure to the vessel until clotting occurred naturally. The mouse was then placed in a custom made holder. The liver was then exposed and directly adhered to a cover slip that was secured in the holder. Once stable the preparation was transferred to a Fluoview FVMPE-RS multiphoton microscope system (Olympus) equipped with a XLPLN25XWMP2 objective (25x; NA1.05; water immersion; 2mm working distance). The laser was 860 nm with a tissue penetration of approximately 50 *μ*m. Coloring depended on the content of the experiment: for experiments with 2 colors, we used violet and green (BA495-540nm + BA410-455nm), while for experiments with 3 colors, we used red, violet, and green (BA575-645nm + BA495-540nm + BA410-455nm). For analysis of the motility of the sporozoites and liver-localized CD8 T cells, we had a tissue penetration depth of approximately 50 *μ*m in a z-stack (2 *μ*m/slice), typically acquired using a standard galvano-scanner at a rate between 0.5 to 1 stacks of 23 images/slices per minute depending on the movie.

Imaging volumes were chosen based on the presence of a parasite; since the Olympus software generates little background autofluorescence and the parasites do not move, this process was dependable. The choice of which parasites to image was contingent upon their appearing in clear, healthy sections of liver tissue, and the proximity of T cells at the time of imaging. Each parasite chosen for imaging required unimpeded blood flow in surrounding sinusoids, bright autofluorescence, and clear surrounding structures. In addition, the parasites chosen for imaging depended on what information we wanted. For the 0 hour timepoint, parasites were selected if they had healthy tissue and had no T cells within a 40 *μ*m radius of the parasite. This allowed for an analysis of attraction of T cells with no primary signaling from an infected cell. At 2 hours post infection, parasites were selected if they had healthy tissue and had ≥ 1 T cells within a 40 *μ*m radius of the parasite. This allowed for an analysis of attraction of T cells with at least 1 primary cell signaling from an infected hepatocyte. We used the 40 *μ*m radius because it has been used to represent the average width of a standard murine hepatocyte^11^.

### Imaging data processing details

We analyzed raw imaging data using the Imaris x64 software (Bitplane) v9.2. We performed the tracking of individual cells in a z-stack using the “Spots” function in surpass mode in Imaris. Exten-sive manual adjustments were subsequently done to ensure the accuracy of the tracks. The detection of individual cells relied upon their relative fluorescence intensity and size (diameter ≥ 9 *μ*m), with a max gap size of 3. We used an autoregressive motion algorithm with background subtraction, tracking enabled, fill gap enabled, and no region growing in Imaris. Since cells are not all in the imaging volume at all times, we used a variable track duration. The detection of fluorescence around each spot was manually adjusted in Imaris prior to tracking cells to reduce background detection at each time point, ensuring the clear distinction of cell versus autofluorescence by the algorithm. Two Imaris files are provided at https://doi.org/10.5281/zenodo.5715658, with one file for the unclustered/small clustered dataset and one file for the large clustered dataset.

### Datasets

1. Dataset #1: unclustered/small clustered data (OT1 T cells in Pb-CS^5M^-infected B6 mice). No or few T cells were near the parasite.
2. Dataset #2: large clustered data (OT1 T cells in Pb-CS^5M^-infected B6 mice). Several T cells were found near the parasite (i.e., with T cell cluster).
3. Dataset #3: no parasite data (OT1 T cells in naive/uninfected B6 mice).
4. Dataset #4: Paris data (PyTCR cells in Py-infected Balb/c mice).
5. Dataset #5: co-clustered (“Kelemen”) data (PyTCR and OT1 cells in Py-infected CB6 mice).

### Metrics

We define four metrics to measure T cell attraction towards the parasite (Figure S3).

1. Metric 1: the angle metric.
  a. For a movement of a cell between timepoints, the angle of movement is defined as the angle between two vectors: the vector from the cell’s position before it moves to the parasite and the vector from the cell’s position before it moves to the cell’s position after it moves.
  b. An acute angle corresponds to “getting closer” and an obtuse angle corresponds to “getting farther”. The probability of randomly getting closer is 0.5.
  c. To test, we associate a Bernoulli distribution with probability 0.5 with each movement, then sum up the distributions for all the movements to get a binomial distribution. Our null distribution is then a binomial distribution with *n* = the number of movements and *p* = 0.5.
2. Metric 2: the improved change of distance metric.
  a. For a movement of a cell between timepoints, the change of distance is defined as the distance from the cell to the parasite after the cell moves minus the distance from the cell to the parasite before the cell moves.

**Figure S1:**
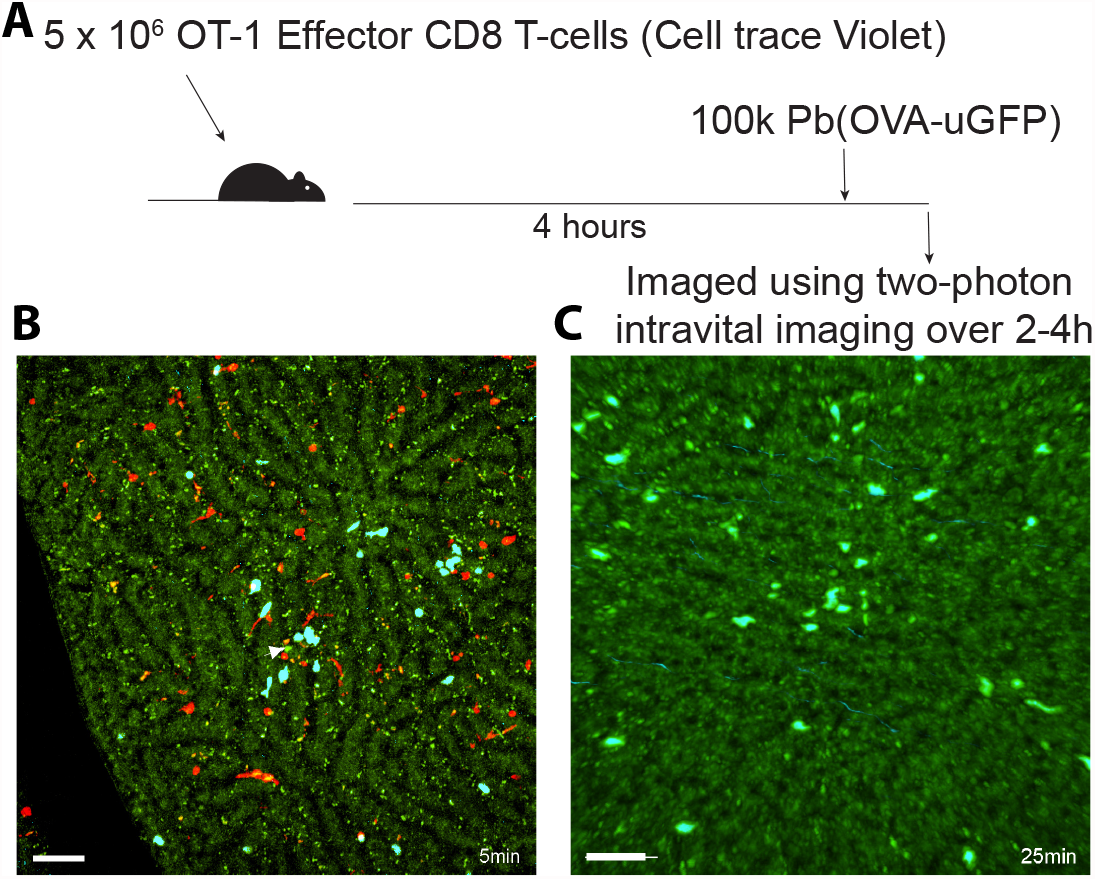
We investigate T cell dynamics in the liver following infection with *Plasmodium* sporozoites. Four hours after i.v. cell transfer of 5 ×10^6^ cell trace Violet labelled OT1 CD8 T cells (blue), mice were injected with 1 × 10^5^ *Plasmodium berghei* Pb(OVA-uGFP) and immediately prepared for intravital imaging (panel A). We imaged the mice using a two-photon standard galvanometer scanner in order to acquire a 50 *μ*m deep Z-stack. We show representative images of small clustering (panel B) and large clustering (panel C) image acquisitions. Representative videos of small clustering (movieS1) and large clustering (movieS2) are in the supplemental materials, as well as the Imaris files at https://doi.org/10.5281/zenodo.5715658. The green sporozoite is highlighted with a white arrow in both images. The scale bars are 20 *μ*m. The numbers overlaid at the bottom right of each image are the timestamps in minutes.

**Figure S2:**
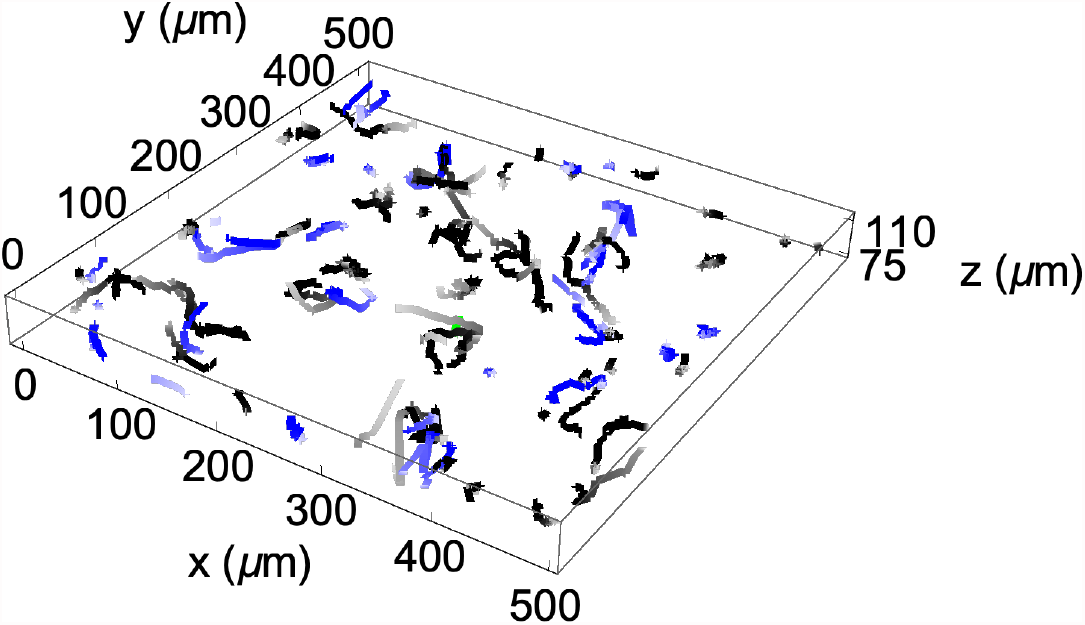
Display of CD8 T cell movements in the liver over time from one unclustered/small clustered dataset. Blue tracks are *Plasmodium*-specific T cells (OT1) and black are nonspecific T cells (P14), with the parasite shown in green (with coordinates (290, 210, 76)). Tracks are shown over time, with early timepoints lighter in color than later timepoints. The data can be explored in greater detail in an interactive Unity environment at https://viktorzenkov.github.io/measuringAttraction.
  b. A negative change of distance corresponds to “getting closer” and a positive change of distance corresponds to “getting farther”. The probability of randomly getting closer is 1*/*2 − *r/*4*x* (Figure S4).
  c. To test, we associate a Bernoulli distribution with *p* = 1*/*2 − *r/*4*x* with each movement, then sum up the distributions for all the movements to get a Poisson binomial distribution, and this is our null distribution.
3. Metric 3: the angle distribution metric.
  a. The angle distribution metric uses the angle of movement from Metric 1 and the angle version of the von Mises-Fisher distribution (described later).
  b. Taking a dataset of angles, we use a log-likelihood test to determine the concentration parameter for the angle version of the von Mises-Fisher distribution most likely to generate the directions corresponding to the angles.
  c. If this distribution’s test statistic is significantly different from a random test statistic, then that corresponds to biased movement, and if it is not significantly different then that corresponds to unbiased movement. If a cell is biased with *κ*_*a*_ > 0, then the cell is considered attracted, and if a cell is biased with *κ*_*a*_ < 0, then the cell is considered repulsed.
  d. A followup test uses the multinomial distribution. For a set of cells, each of which has had its bias ascertained as attracted, repulsed, or unbiased, we set our null distribution to a multinomial distribution with probabilities set to the proportion of cells detected as unbiased, *p*_*u*_, and the other probabilities each set to^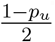^. This takes into account the proportion of cells that are unbiased while still allowing a test between attracted and repulsed quantities.
4. Metric 4: average angle metric.
  a. The average angle metric uses the angle of movement from Metric 1.
  b. To test, we use a Student T test to compare the mean of the angles for a cell to 90 degrees, which represents randomness.

### Statistical/computational analyses

#### A randomly moving cell is more likely to have its distance from a parasite increase

There have been few studies that rigorously address the question of how to detect attraction of a moving lymphocyte towards an infection site. Commonly, two metrics or their variations have been used: the angle-based metric and the distance-based metric^12,16,19^. According to these metrics, cells moving towards the parasite exhibit acute angles to the parasite and reduction in distance to the parasite, respectively (Figure S3A&B). The natural assumption is that randomly moving (unbiased) cells should exhibit a similar number of movements towards and away from infection. However, as far as we know this assumption has not been thoroughly explored.

To evaluate the potential bias of T cells towards the infection site, we performed stochastic simulations in which T cells searched for the infection. Simulations showed that while the angle metric is not biased (i.e., for randomly moving T cells, there are similar proportions of acute and obtuse angles to infection), the distance metric is biased (i.e., there are more movements away from the infection than towards the infection). (Note that this usage of the word “bias” refers to a skewed result, unlike our other usage, which refers to cell movement being attracted or repulsed.) Even before performing simulations, graphing movement possibilities of T cells with respect to the infection site revealed that for a finite distance between the cell and the infection, there are more chances for the T cell to move away than to get closer (Figure S4). Defining *r* as the distance that the cell moves and *x* as the distance from the cell to the parasite, the probability that the distance decreases for a T cell movement is 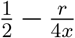. The subset of the surface of the *r*-sphere located inside the *x*-sphere are the positions for which the cell’s distance to the parasite gets smaller, and the subset of the surface of the *r*-sphere located outside the *x*-sphere are the positions for which the cell’s distance to the parasite gets larger. Note that if *r* > 2*x*, causing the fraction to be less than 0, then the cell moves so far that none of its potential destinations are closer to the parasite than its current location, which means the probability that the distance gets smaller is 0 (Figure S4).

**Figure S3:**
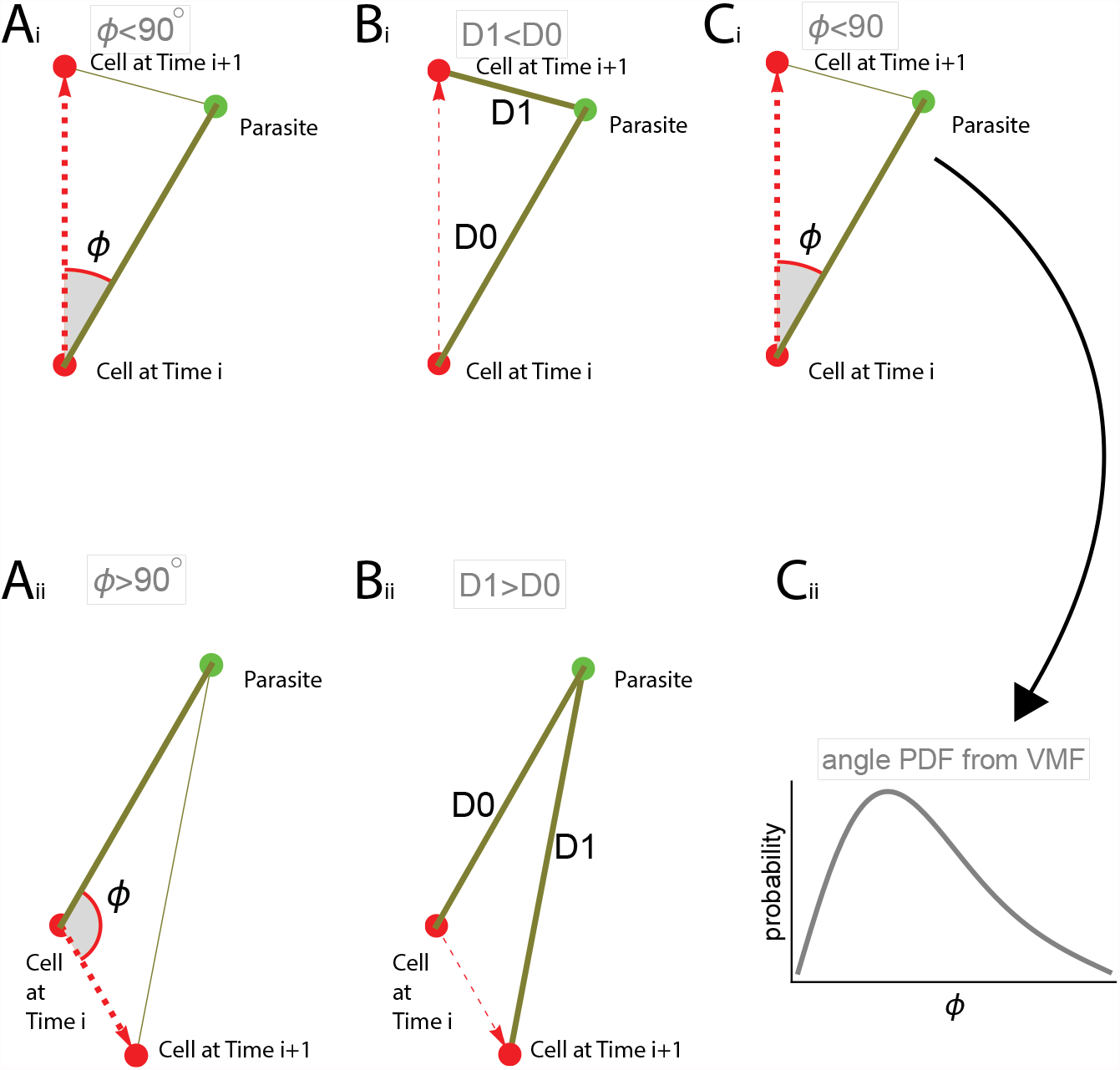
Four metrics were used to test for attraction of CD8 T cells towards the infection site. Panels A: the angle metric is defined as the angle between the vector from the cell to the parasite and the vector formed by the cell’s movement. An acute angle corresponds to “getting closer” (Ai) and obtuse corresponds to “getting farther” (Aii). The fourth metric, the “average angle” metric, uses these same angles. Panels B: the distance metric is defined as the change between the distances from the cell to the parasite before and after the cell moves. A negative change of distance corresponds to “getting closer” (Bi) and positive corresponds to “getting farther” (Bii). Panels C: the angle distribution metric takes into account the actual values of angles and compares these values with the von Mises-Fisher distribution (eqn. (1)) leading to an estimate of the concentration parameter *κ*_*a*_ (a measure of attraction). If *κ*_*a*_ > 0 and this distribution’s test statistic is significantly different from a random test statistic (*p* < 0.01), we consider the cell to have “attraction”. If the test statistic is not significantly different, then we consider the cell to not have bias. If *κ*_*a*_ < 0 and the test statistic is significantly different, then we consider the cell to be repulsed.

**Figure S4:**
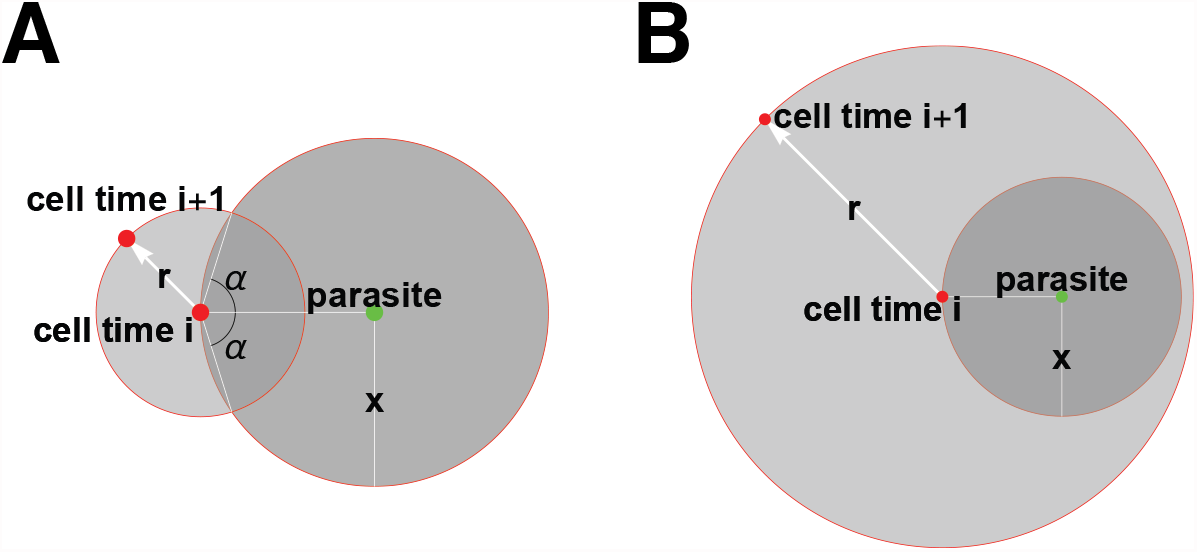
This graphic demonstrates the bias in the distance metric’s measure of attraction to the infection site. Based on the volumes of the areas the cell may move to, the probability that a T cell moves closer to the parasite is lower than the probability that the T cell moves away from the parasite. With *r* defining the distance that the T cell moves and *x* defining the distance between the cell and the parasite, the probability that the distance between a randomly moving T cell and the parasite declines with the movement is *P* (Δ*D* < 0) = 1*/*2 − *r/*(4*x*) for *r* < *x* (Panel A) and is *P* (Δ*D* < 0) = 0 if *r* > *x* (Panel B).

Thus, the distance metric is biased in the sense that movements have a less than 50% probability of getting closer to the parasite, which means our null distribution cannot define randomness as 50%. To form a corrected test, we utilized a Poisson binomial distribution as our null distribution, which is a sum of Bernoulli distributions with different probabilities (in our case, a Bernoulli for each movement). The probability that *k* movements (out of *n*) are towards the parasite is then:

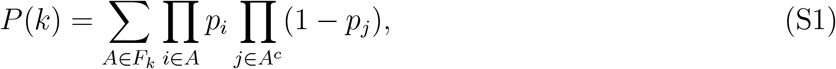

where *F*_*k*_ is the set of all subsets of *k* integers that can be selected from 1, 2 … *n* and *A*^*c*^ is the complement of *A, p*_*i*_ = 1*/*2 − *r*_*i*_*/*(4*x*_*i*_) and *r*_*i*_ is the length of the *i*^*th*^ cell movement and *x*_*i*_ is the distance between the T cell and the parasite before the *i*^*th*^ movement (Figure S4). With this correction, the angle-based and distance-based test results are numerically indistinguishable.

#### The von Mises-Fisher distribution provides a way to model and measure attraction

One of the limitations of the angle- and distance-based metrics is that the actual value of the angle/distance is ignored and converted into a boolean value, thus potentially reducing the power of detecting bias in cell movement. To retain the actual values of the angles in a test, we used the von Mises-Fisher (vMF) distribution. The vMF distribution is a natural extension of the von Mises distribution used in ecology, which is restricted to describing bias in 2D^20^.

The vMF distribution chooses an *n*-dimensional (we use *n* = 3) vector given a direction vector *μ* and a concentration parameter *κ*^25,26^. Sampling from the vMF distribution provides a vector chosen pseudorandomly with a bias toward the given vector whose strength is dependent on *κ*, with *κ* → 0 having no bias, *κ* > 0 having positive bias (attraction), and *κ* < 0 having negative bias (repulsion). The probability density function for a vector *χ* in the vMF distribution in 3D is

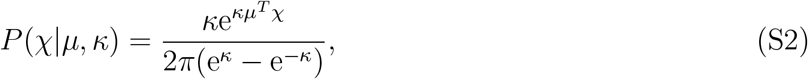

where *μ* is the direction vector toward which there is bias and |*μ*| = |*χ*| = 1. To calculate the angle *ϕ* between vectors *χ* and *μ*, we let *μ* = (0, 0, 1) and *χ* = (*x, y, z*). Given that |*χ*| = 1, the angle *ϕ* then only depends on the value of *z* and can be calculated from the relationship cos(*ϕ*) = *z*. With that transformation and utilizing Mathematica we found the following probability density function for the angle *ϕ*:

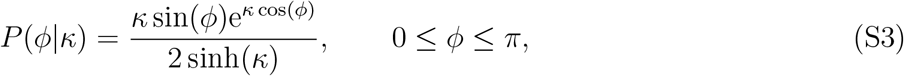

where *ϕ* is the angle between the vector to the attraction point and the cell movement vector. Note that in a case with no attraction, 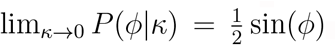. Because the magnitude of the concentration parameter is not intuitive, another useful parameter is the fraction of acute angles towards the attraction site. This fraction is found by integrating the vMF distribution

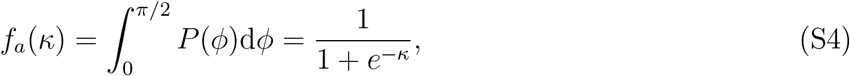

where *f*_*a*_(0) = 1*/*2 and *f*_*a*_(∞) = 1.

#### Estimating the concentration parameter of the vMF distribution

First, we sum the logs of the probabilities of getting each experimentally determined angle to the parasite *ϕ*_*i*_ from the vMF angle PDF (eqn. (1)) to get the negative log-likelihood function

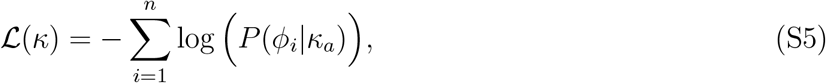

where *n* is the number of movements in the data. The maximum likelihood estimate *κ*_*a*_ is found by minimizing eqn. (S5) (e.g., in Mathematica using the function Maximize on the inverse of the expression). Whether the found estimate 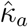 is different from 0 is determined by using a likelihood ratio test (LRT) to compare 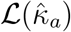 and 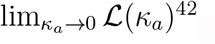.

It is straightforward to use a mixture of vMF distributions to explain angle distribution data. In particular, we found that while a single vMF distribution describes the distribution of angles to the parasite reasonably well, there is an over-representation of 90^*°*^ angles in the data (e.g., Figure S12). Therefore, in this case we fit the following mixture distribution to the data

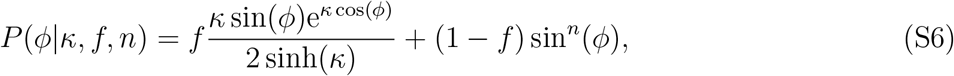

where *f* is the fraction of cells that are biased towards the infection, sin(*ϕ*) is proportional to a vMF distribution with no bias, and *n* is used to explain the over-representation of angles at 90^*°*^. Improvements of this model fit over the simpler model, based on a single vMF distribution, are tested using a likelihood ratio test.

#### Modeling T cell attraction to the parasite

We use the vMF distribution to model T cell attraction to the parasite by using angles to infection as the set of angles, and we define *κ*_*a*_ as the concentration parameter determining the strength of this attraction.

#### Modeling a correlated (persistent) random walk

Previous studies of cell movement employed various methods to simulate a persistent random walk, such as assuming a normal distribution of turning angles with mean 0 and some variance^41,43^. Other studies used the Ornstein-Uhlenbeck model to simulate persistent random walks^29,44,45^, which we also model in a later section. Alternatively, we use the vMF distribution to model persistent random walks using the concentration parameter *κ*_*t*_ for turning angles. In this model, turning angles are sampled from a vMF distribution with concentration *κ*_*t*_. There is a direct relationship between the average turning angle 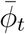 of a cell moving in a persistent random walk and the concentration *κ*_*t*_:

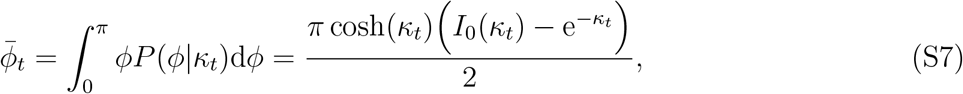

where *I*_0_(*κ*) is the Bessel function of the first kind of order 0.

#### The von Mises-Fisher distribution-based test may also be improved to eliminate bias occurring in a correlated random walk

The vMF distribution-based metric is not biased if cells move randomly with respect to a cell’s previous movement. However, in many situations T cells move with a persistent random walk^17^: cells move in a direction positively correlated with the previous movement direction. To investigate if a persistent random walk may introduce bias in estimating attraction towards the infection site, we performed a set of simulations in which we describe cells’ movement by our vMF distribution: the angle *ϕ* determines the cell’s turning angle and the concentration parameter *κ*_*t*_ indicates the degree of the cell’s persistence in the random walk. Analyses suggested that a persistent random walk can cause some cells to appear attracted to the infection site (or repulsed from the infection site) even though there was no actual attraction or repulsion (Figure S6).

Interestingly, we found that the fraction of cells detected as biased in simulations depended on the degree of walk persistence and the size of the volume in which the movement of T cells was considered, but in different manners. For low values of *κ*_*t*_ (low persistence), the fractions of cells detected as attracted and repulsed were similar (both low). For higher values of *κ*_*t*_ (high persistence), the fraction of cells detected as attracted was not impacted by the size of the volume, but the fraction of cells detected as repulsed was impacted: in all simulations with the maximum simulated persistence *κ*_*t*_ = 2, about 6% of cells were detected as attracted, while the percent of cells detected as repulsed ranged from about 6% to about 11% as the volume increased from a small box (Figure S6Aii) to an infinite volume (Figure S6Cii). This is not surprising, however, because cells that are detected as repulsed tend to move away from the parasite and are more likely to exit smaller imaging volumes (as in the smaller box in Figure S6Ai).

An interesting but perhaps unrealistic imaging experiment would be a situation when there is no “imaging” box. According to the simulations, this scenario results in the greatest disparity between the fractions of cells detected as attracted and repulsed. This phenomenon occurs for cells with greater persistence because of the artifact that cells moving away from the parasite tend to continue to move away, while cells moving towards the parasite tend to pass the parasite and then also move away. We could not derive an analytical expression to correct for the bias in detecting attraction of cells moving in a correlated random walk. However, by simulating the movement of cells using the cell’s initial position, movement length distribution, and turning angles as observed in the data, but with no attraction, it is possible to determine the bias of T cell movement towards the infection site in the absence of actual attraction,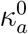. We thus used this estimate for 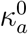 in our test of whether the detected bias *κ*_*a*_ in actual data was significantly different from 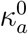.

**Figure S5:**
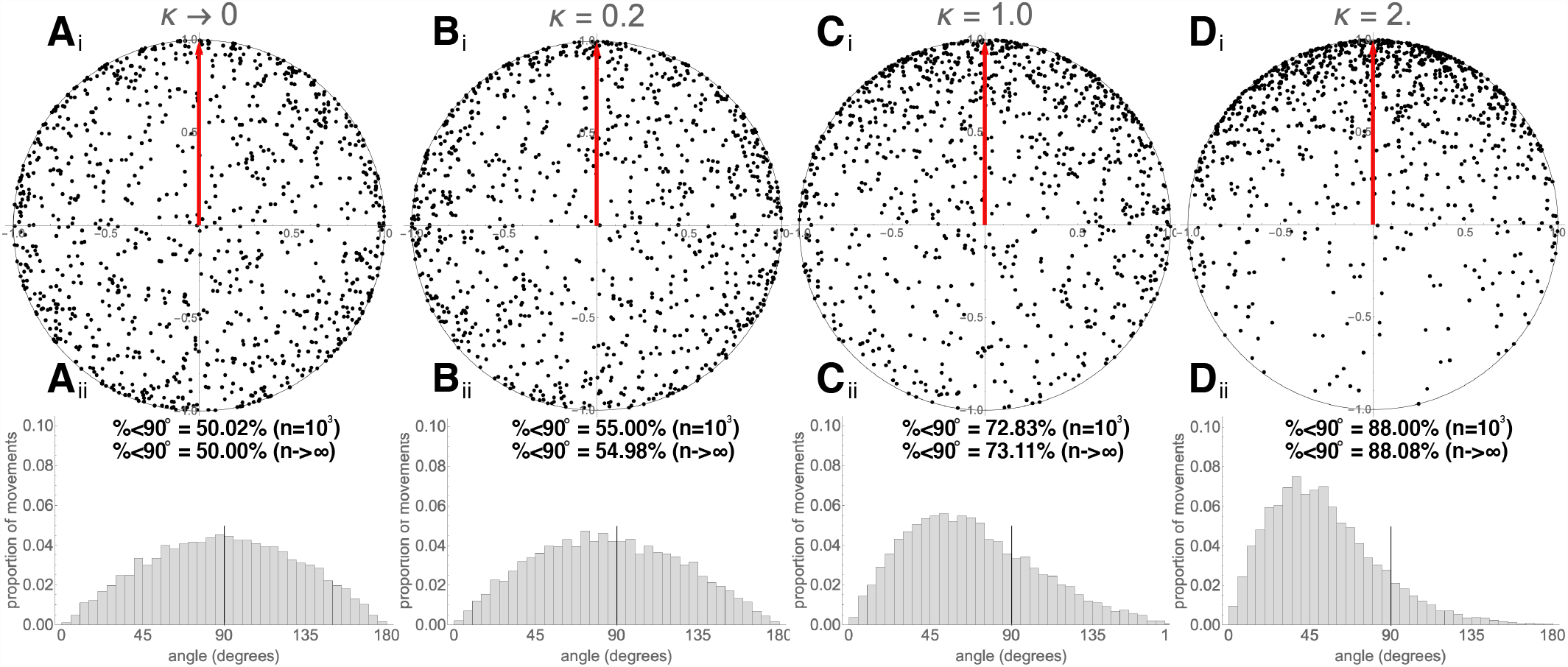
Illustration of the random vectors generated using the von Mises-Fisher (vMF) distribution. The vMF distribution generates direction vectors with preference toward a given direction, with the strength of preference determined by the concentration parameter *κ*. We sampled 1000 directions from a vMF distribution for several values of *κ* (noted on individual panels) and plotted those in a 2D image by removing the z-coordinates (panels i). We also show histograms of the actual angles between the movement vector and the vector to the parasite (panels ii, angle metric defined in Figure S3). In panels ii we also show the percents of acute angles generated for the *n* = 1000 pseudorandomly sampled vectors which were calculated using eqn. (S4).

#### How to create stochastic simulations based on the vMF distribution

We simulated a moving cell by generating a set of movements, each of which is defined by a distance traversed and a direction vector of the cell’s movement. To generate the movement distances, we used Mathematica to estimate the best distribution to fit the distances from a real dataset, which resulted in the Pareto Type IV (generalized Pareto, GP) distribution:

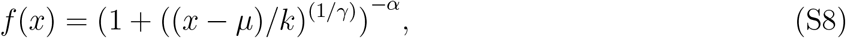

with the following best fit parameters: minimum value *k* = 7, location parameter *μ* = 0, scale parameter *α* = 2.7, and inequality parameter *γ* = 0.68, assuming that these movements occur in 2 min intervals. These parameters were found by fitting the GP distribution to movement data from datasets #4 and #5 pooled together, using the function `FindDistributionParameters` on a `ParetoDistribution` in Mathematica. Of note, we also found that the GP best fits the movements of malaria-specific CD8 T cells in the absence of a malaria infection^39^.

To simulate attracted movement of T cells towards a parasite, each direction was chosen from the vMF distribution with the angle between T cell movement and the parasite given in eqn. (1). The concentration *κ*_*a*_ was chosen based on the desired amount of attraction (*κ*_*a*_ > 0 corresponds to attraction, *κ*_*a*_ = 0 corresponds to no bias, and *κ*_*a*_ < 0 corresponds to repulsion). To create the desired vector, we first chose a vector with bias toward direction {0,0,1}, which simplifies the process to choosing *x* and *y* pseudorandomly from a normal distribution *N* (0, 1) and choosing *z* based on the von Mises-Fisher distribution, using

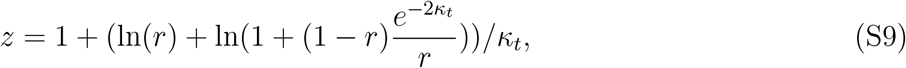

where *r* is chosen uniformly between 0 and 1, exclusively^46^. Then we weighted *x* and *y* to place the chosen vector on the unit sphere, and used a rotation transform to adjust the generated vector with respect to the desired bias direction.

To simulate a correlated (persistent) random walk, each movement direction was chosen from a vMF distribution with a vector input of the vector from the cell’s previous position to the cell’s current position. The concentration *κ*_*t*_ was chosen based on the desired amount of movement persistence, with *κ*_*t*_ > 0 corresponding to persistence, *κ*_*t*_ = 0 corresponding to randomness, and *κ*_*a*_ < 0 corresponding to the cell turning back on itself. For a constant parasite position, a sequence of movements was simulated consecutively, resulting in a list of cell positions similar to those from the experimental datasets.

We also simulated a walk with both attraction to a parasite and persistence by choosing two directions with bias as described previously and summing each pair of directions to find a direction with bias influenced by both the parasite and movement persistence. This framework can easily be extended to more than two biases in a similar way.

#### How to create stochastic simulations based on the Ornstein-Uhlenbeck process

The Ornstein-Uhlenbeck (OU) process is a stochastic process that generates a correlated (persistent) random walk^16,45^. The OU process chooses each position to move to using the following formula:

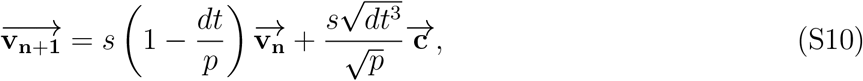

where 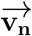 is the *n*^th^ position of a cell, *s* is the speed, *p* is the persistence time, *dt* is the timestep in simulations, and 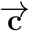 is a pseudorandomly chosen vector to introduce randomness. In this model, *p* determines the timescale at which movements are correlated (i.e., the timescale of the persistence of the random walk). We adjusted the OU process to simulate cells with attraction to a specific point in space, rather than relative to the previous movement vector, using the following formula:

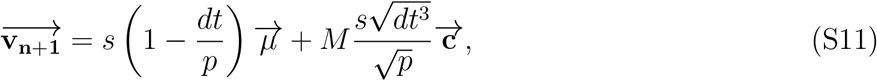

where 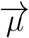 is a vector from the cell’s current position to the parasite, *M* is a multiplier that increases the effect of the randomness, and other parameters are the same as the persistent OU model (eqn. (S10)). In this model, the strength of attraction to the parasite is determined by *p*, with larger values determining stronger attraction, and *M* negates the impact of such attraction by introducing higher degree of randomness in the movements. For each analyses, we simulate cell’s positions at a regular interval *dt* which is typically small (e.g., 1 second), and then we sample every 100th position to gain simulated data comparable to real cells (one position per 100 seconds). We typically chose the following parameters to simulate walks with attraction towards the parasite: *s* = 1.2*μ*m*/s, p* = 1.1*s* to 10.1*s, dt* = 1*s, M* = 500. In these simulations, cells have a starting distance of 100*μ*m.

#### The von Mises-Fisher distribution-based metric is the most powerful metric at detecting attraction from the tested alternatives

We performed two sets of stochastic simulations in order to determine which of our metrics is most powerful, i.e., which metric requires the least amount of data to detect attraction, or equivalently, which metric detects more/stronger attraction for the same amount of data. In the first set of simulations, we described movement lengths of T cells by the generalized Pareto distribution and described attraction towards the infection site by the vMF distribution with different concentration parameters. In the second set of simulations, we described the movement of T cells using the modified OU process with different persistence times (see eqn. (S11)). We found that independent of the method used to simulate attracted movement for the same number of movements (data points), the vMF distribution-based metric allowed us to detect the smallest bias in movement. Equivalently, for the same degree of attraction, the vMF distribution-based metric required the least data to detect attraction (Figure S7). This demonstrates the stronger sensitivity of the vMF-based metric. We also note that as *κ*_*a*_ approaches 0, the percent of cells detected as attracted decreases, indicating a small false positive rate.

---

**Figure S6:**
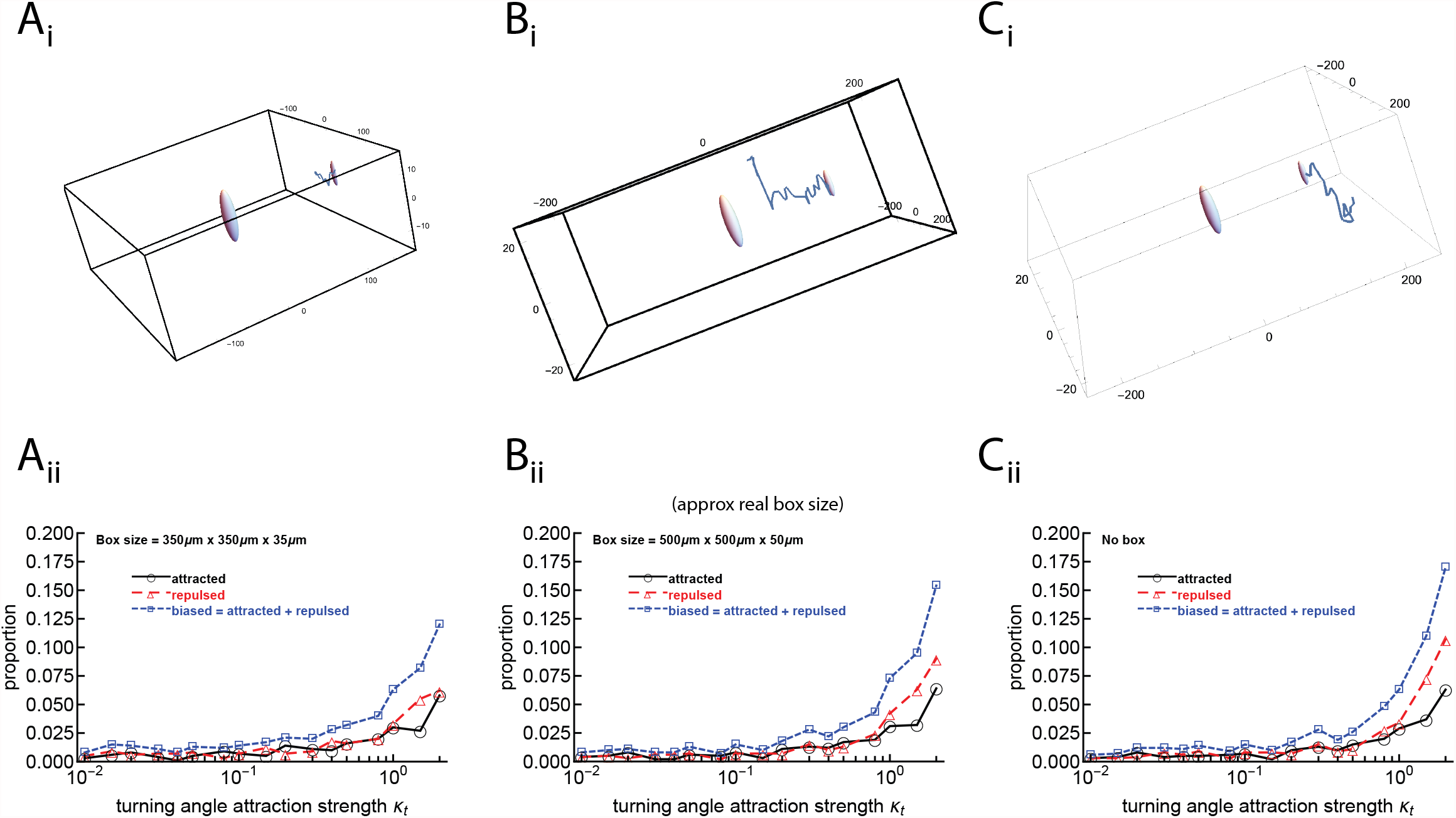
We quantified bias in the angle distribution (vMF distribution-based) metric and found that the number of cells detected as repulsed depends on the size of the imaging volume when the cells are persistent. We run stochastic simulations of T cells moving in a small “boxed” space (350 ×350 ×35 *μ*m, panels A), a regular boxed space as used in intravital imaging experiments (500 ×500 ×50 *μ*m, panels B), and in open space (panels C). The parasite is assumed to be located in the center of the box and T cells start movement at 150 *μ*m from the parasite at coordinates (*x, y, z*) = (150, 0, 0). We assume 50 movements per simulated T cell, with movement lengths described by the Generalized Pareto distribution (eqn. (S8)) with parameters given in Materials and Methods. To simulate a persistent random walk, we assume that T cell turning angles are described by a vMF distribution with constant *κ*_*t*_ determining the bias of a T cell walk (see Materials and Methods for more detail). A cell is assumed to be out of the imaging volume when the cell’s position is outside of the defined imaging box (in A and B) which then results in fewer than 50 moves recorded. For each cell’s track, we determine if the cell is attracted (*κ*_*a*_ > 0), repulsed (*κ*_*a*_ < 0), or unbiased (*κ*_*a*_ = 0) to the parasite’s position by comparing the distribution of angles to the parasite to a vMF distribution (Figure S3) and estimating the concentration *κ*_*a*_. For each concentration determining the persistence of the random walk *κ*_*t*_, we simulate movement of 1000 cells. In Panels i are examples of a cell whose path is cut off by the small box (Ai), a cell detected as attracted (Bi), or a cell detected as repulsed (Ci). Note that in Panels i the figures are not to scale, resulting in the spheres representing the cells being elongated. In these examples, the parasite and cell’s starting position and the cell’s track are shown. Parasites are depicted as a sphere with a radius of 40 *μ*m representing the infected hepatocyte^10,11^.

**Figure S7:**
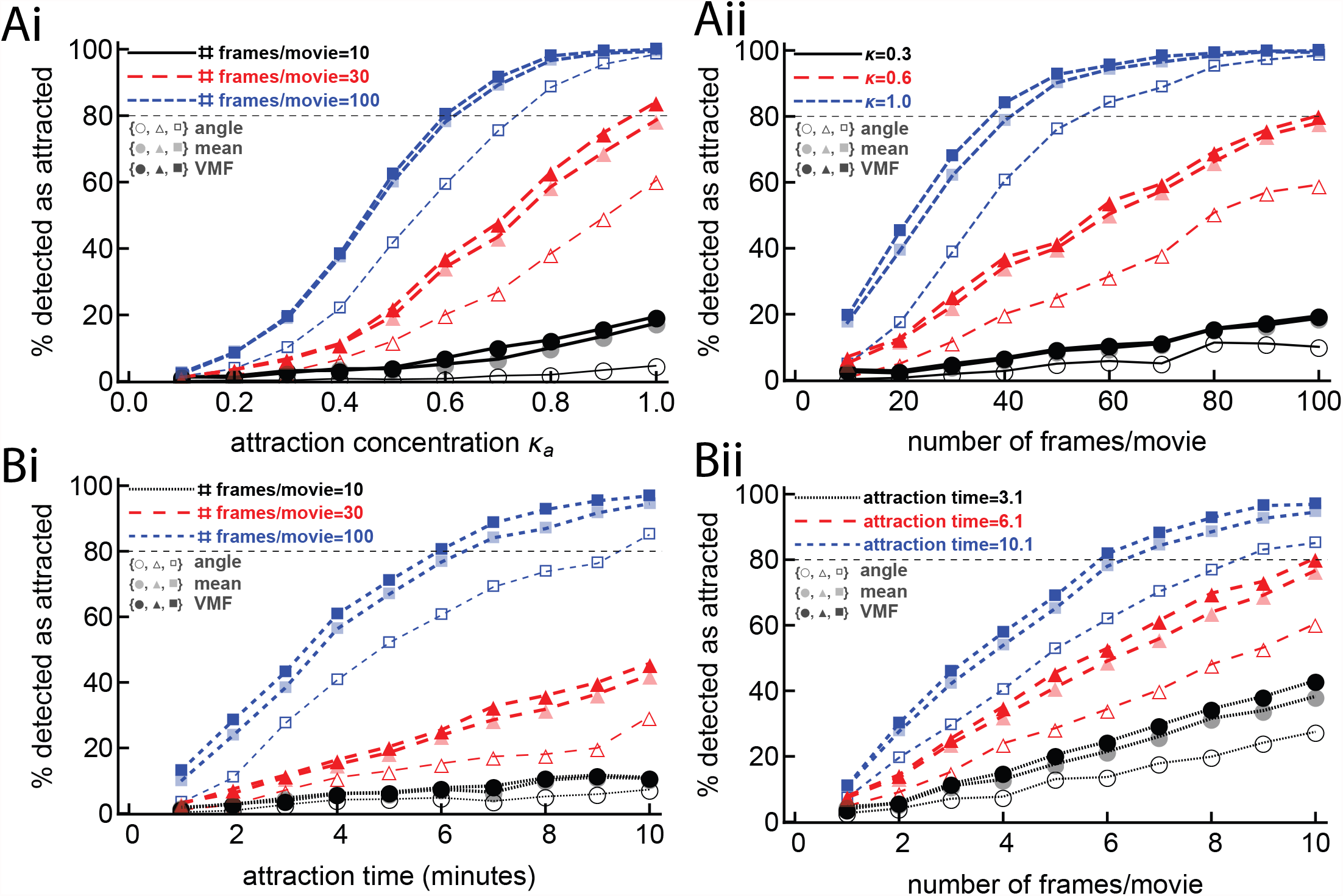
The third metric, based on the vMF distribution, is the most powerful out of the tested metrics (demonstrated by its percent of cells detected as attracted always being higher than the other metrics). We simulated T cell movement with varying degrees of bias towards the parasite for different movement durations using two different frameworks: using vMF distribution-based simulations (panels A and Figure 4) or using the Ornstein-Uhlenbeck (OU) process (panels B and eqn. (S11)). For each simulated trajectory we determined the percent of cells detected as attracted using the angle metric (metric 1, open symbols), the vMF distribution-based metric (metric 3, solid symbols), and the mean angle metric (metric 4, transparent symbols). The distance metric (metric 2) is not shown to avoid clutter and because it performs similarly to metric 1. Cells were detected as attracted if their movement was statistically different from unbiased. In total, we simulated 1000 cells per every set of parameters (see Additional experimental details section for exact values of parameters used in simulations).

**Figure S8:**
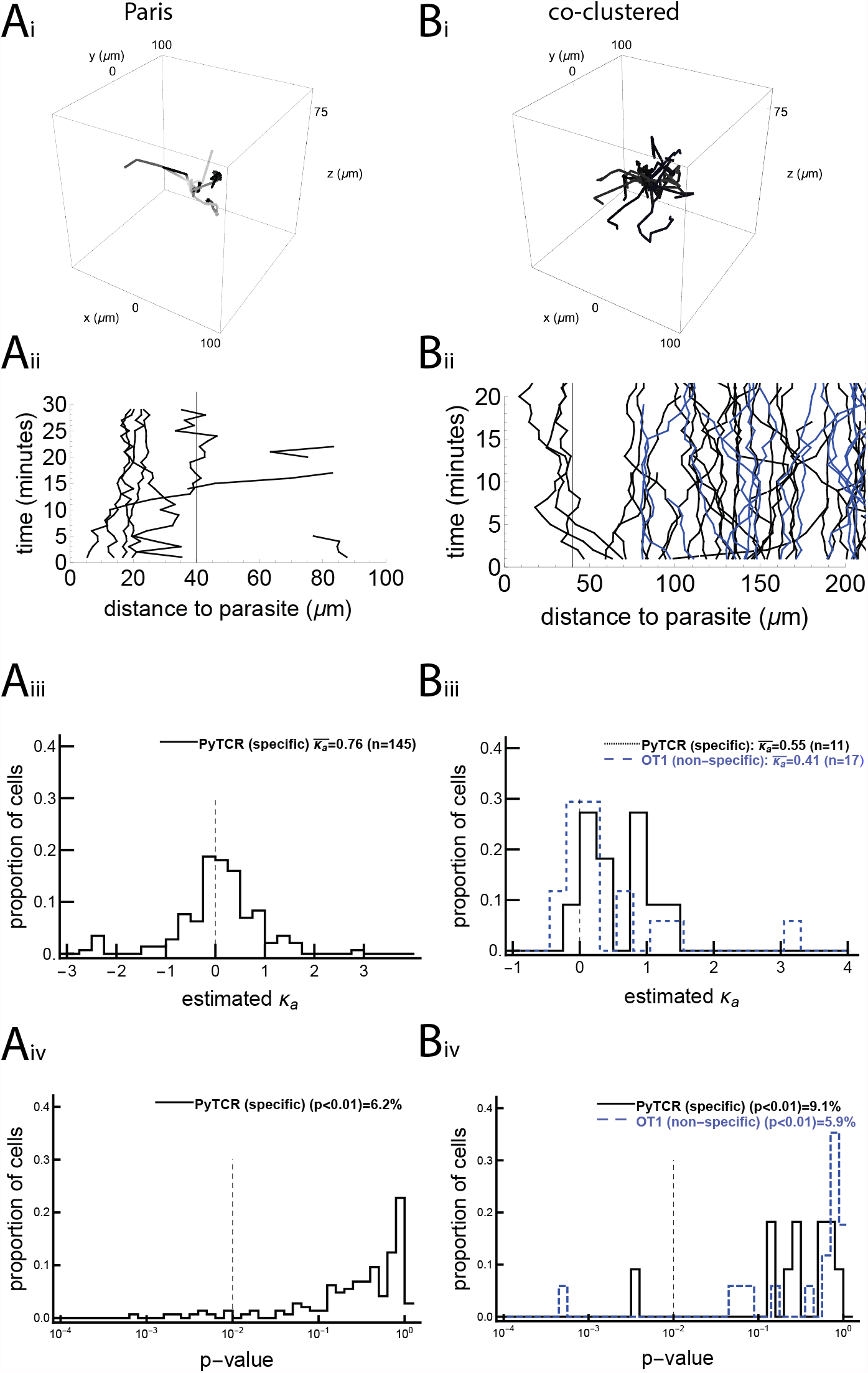
A minority of activated CD8 T cells display movement bias to the liver stage. This analysis was similar to Figure 1, for the previously published (“Paris” and “co-clustered”) data^10,12^. In the Paris data there were no nonspecific cells, but there were four mice with several infection sites each. In the co-clustered data, we imaged CD8 T cells, specific to Py (PyTCR) or specific to a irrelevant antigen (OT1), in one mouse with one Py liver stage. For the Paris data, we detected 5 PyTCR cells as attracted and 4 PyTCR cells as repulsed. For the co-clustered data, we detected 1 PyTCR cells and 1 OT1 cells as attracted and 0 PyTCR cells and 0 OT1 cells as repulsed.

**Figure S9:**
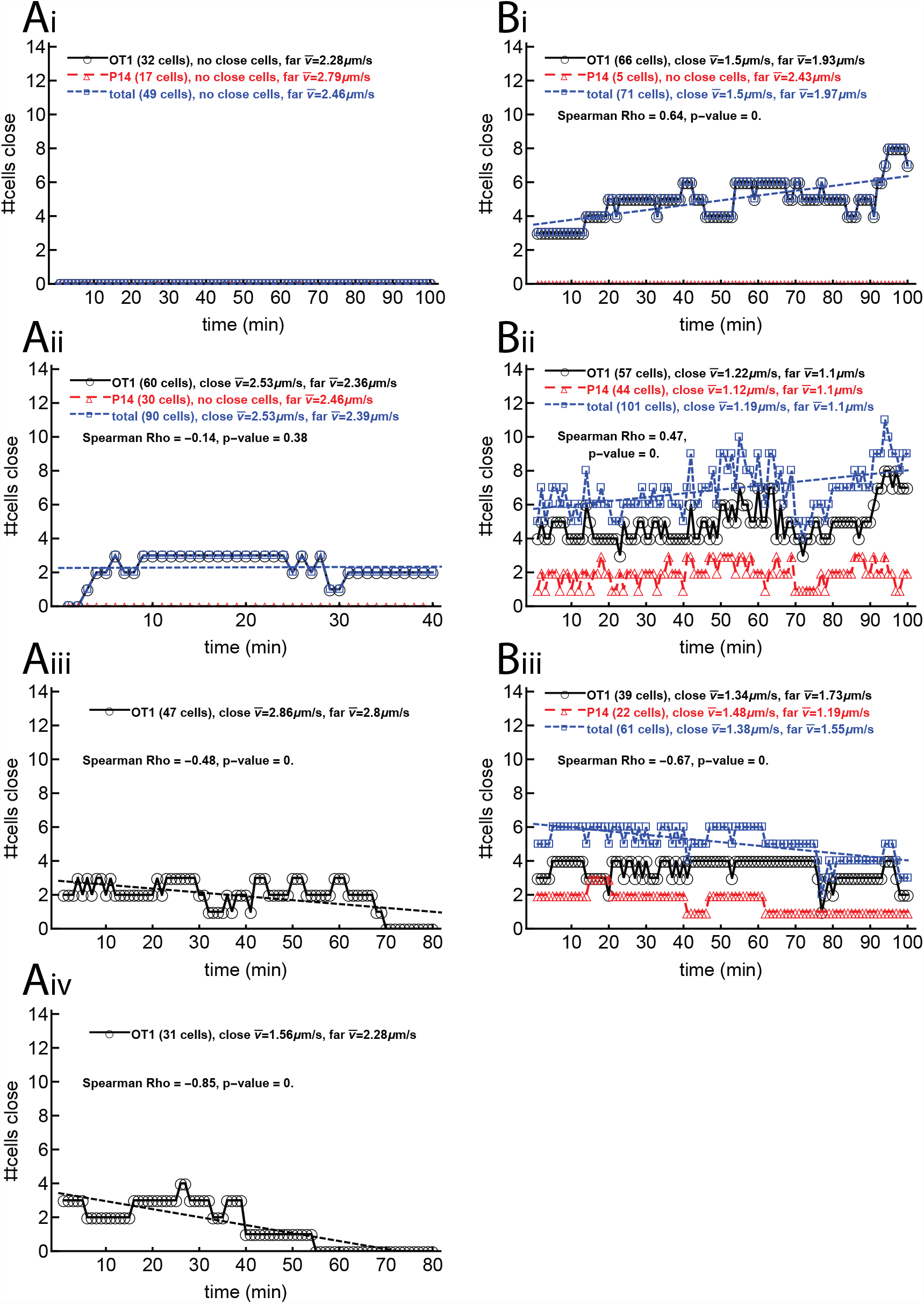
The numbers of cells close to the parasite change more over time for large clustered datasets than for unclustered/small clustered datasets, according to the Spearman correlation test. The size of the cluster (the number of cells close to a parasite) is of interest, because if the cluster size increases, then cells are clearly coming to the parasite and not leaving. Here we show the number of cells close to the parasite (within 40*μ*m from the parasite) at each time point, with one panel for each parasite. The left column is for the unclustered/small clustered dataset, and the right column is for the large clustered dataset.

**Figure S10:**
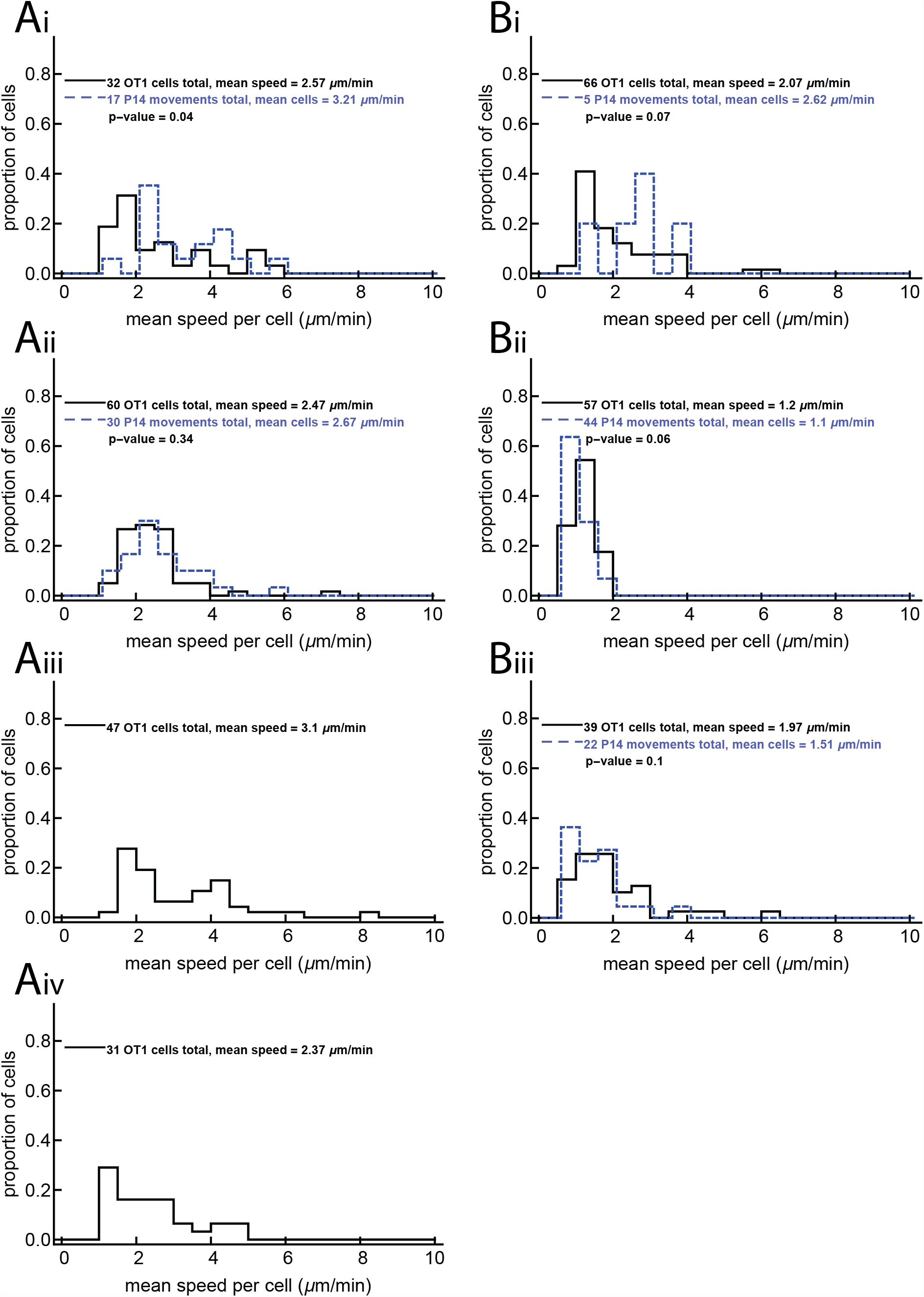
Distribution of average speed per cell in different imaging experiments. Left panels are for the unclustered/small clustered dataset, and right panels are for the large clustered dataset. The p-values come from the Mann Whitney test.

**Figure S11:**
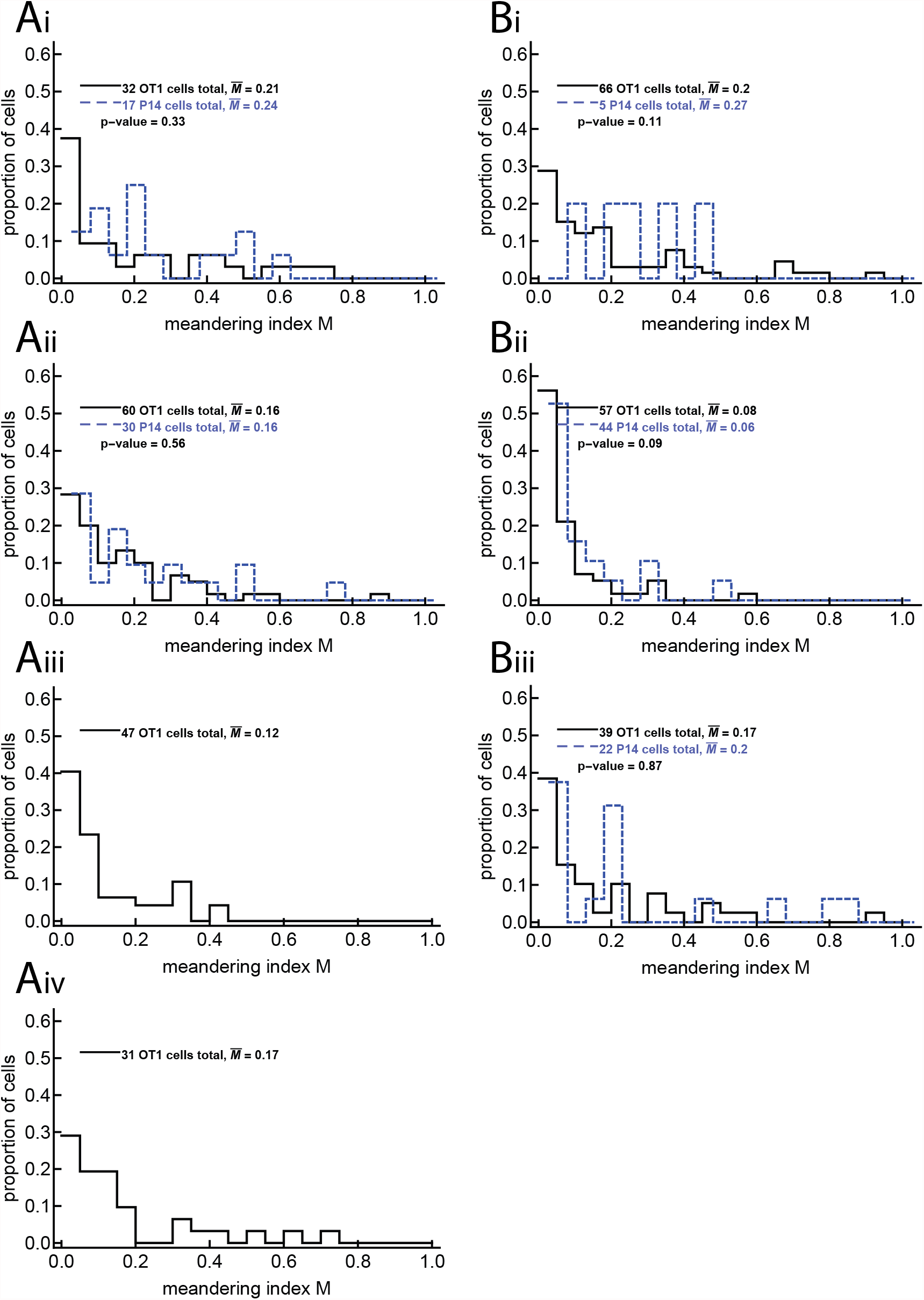
We display the distributions of meandering indices for cells in the experiments with dataset #1 (unclustered/small clustered dataset) and #2 (large cluster dataset). The meandering index *M* is calculated as the distance between the first and last recorded positions of the cell divided by the total length of the cell’s movement in space; it is therefore between 0 and 1, with 0 equivalent to the cell returning to its original position and 1 meaning the cell moved in a straight line. In our real data, the meandering indices are mostly around 0.15 to 0.2, and are identical between corresponding specific and non-specific datasets. The p-values come from a Mann Whitney test.

**Figure S12:**
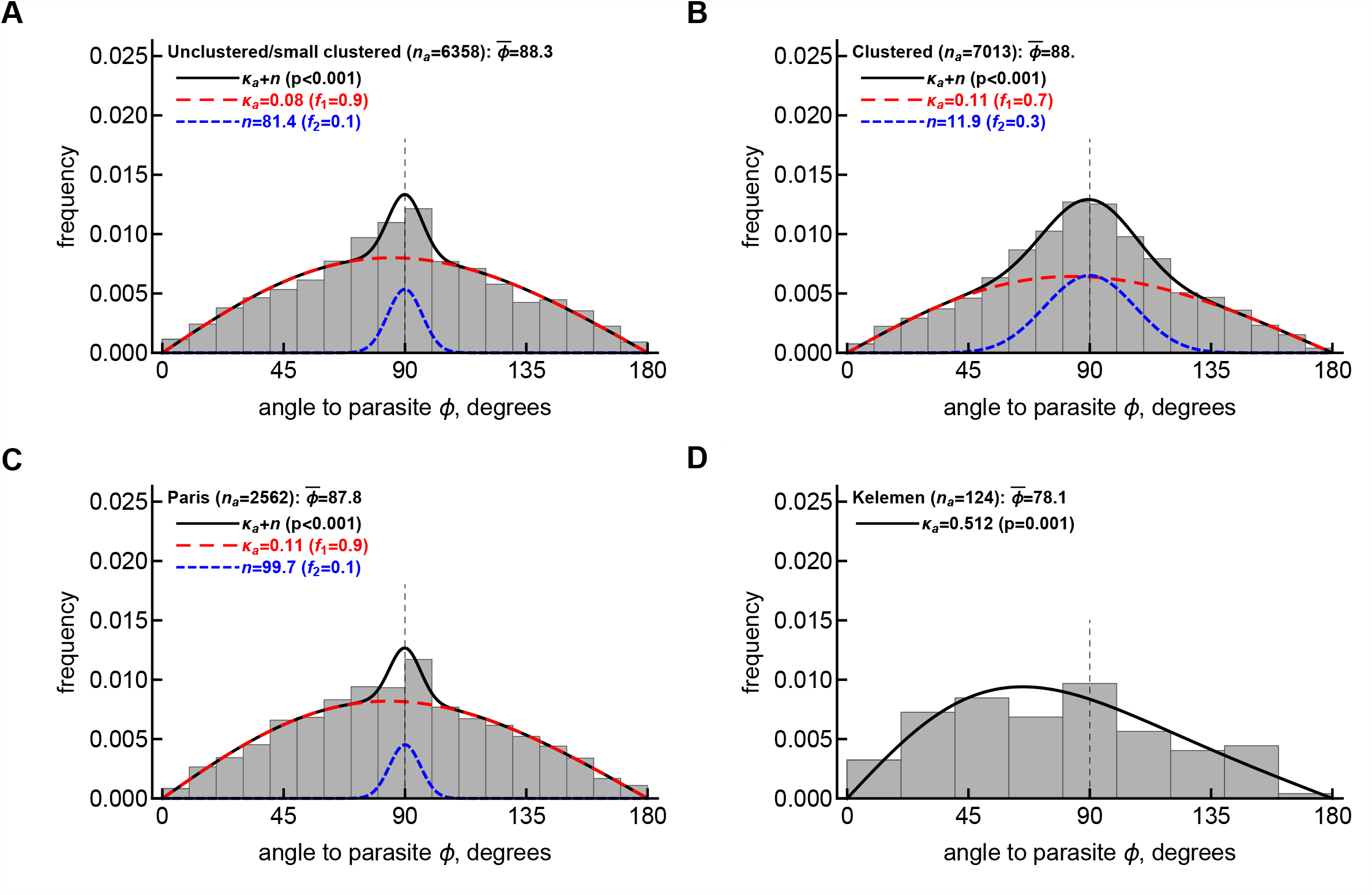
The von Mises-Fisher distribution describes relatively well the distributions of angles to the parasites from four datasets. We show the distributions of angles to the parasite of *Plasmodium*-specific liver-resident CD8 T cells, as outlined in Materials and Methods, for the unclustered/small clustered dataset (A), clustered dataset (B), Paris dataset (C), and co-clustered dataset (D). We fit either a vMF distribution (given in eqn. (1)) or a mixture of distributions (eqn. (S6)) to these data and estimated the concentration parameter *κ*_*a*_ (noted on individual panels). Fits of the model are shown by lines. We compared every fit with the fit of a simpler model (a single vMF distribution fit in panels A-B or a null distribution of angles with *κ*_*a*_ → 0 in panel D) using a likelihood ratio test; resulting p-values are shown on individual panels. For each dataset we also plot the total number of angles in the dataset (*n*_*a*_) and the average angle to the parasite 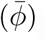.

**Figure S13:**
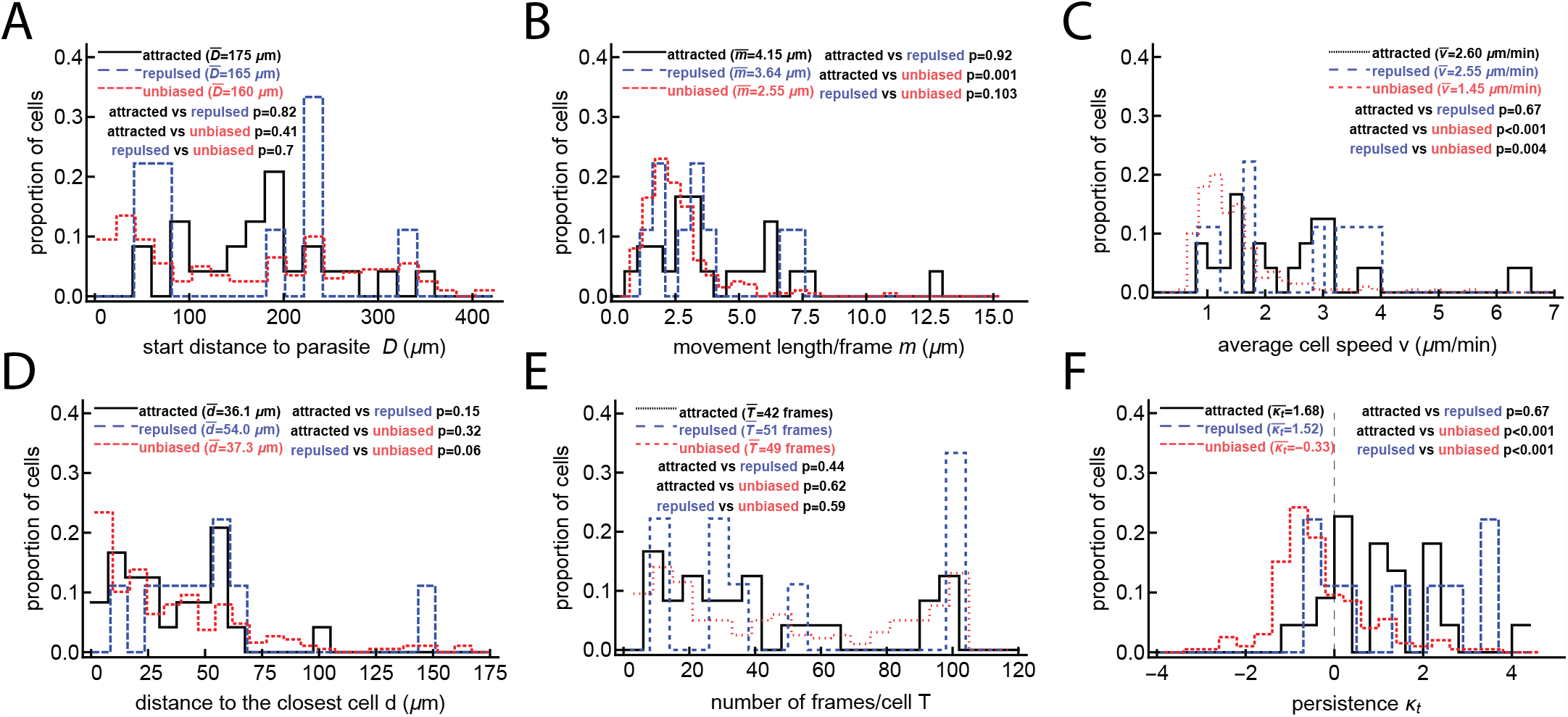
Speed and turning angle both correlate with detection of attraction when several T cells have found the parasite. These are metadata results from tests that use data divided into attracted, repulsed, and unbiased T cells for the large clustered dataset (dataset #2). For more details, see Figure 2.

**Figure S14:**
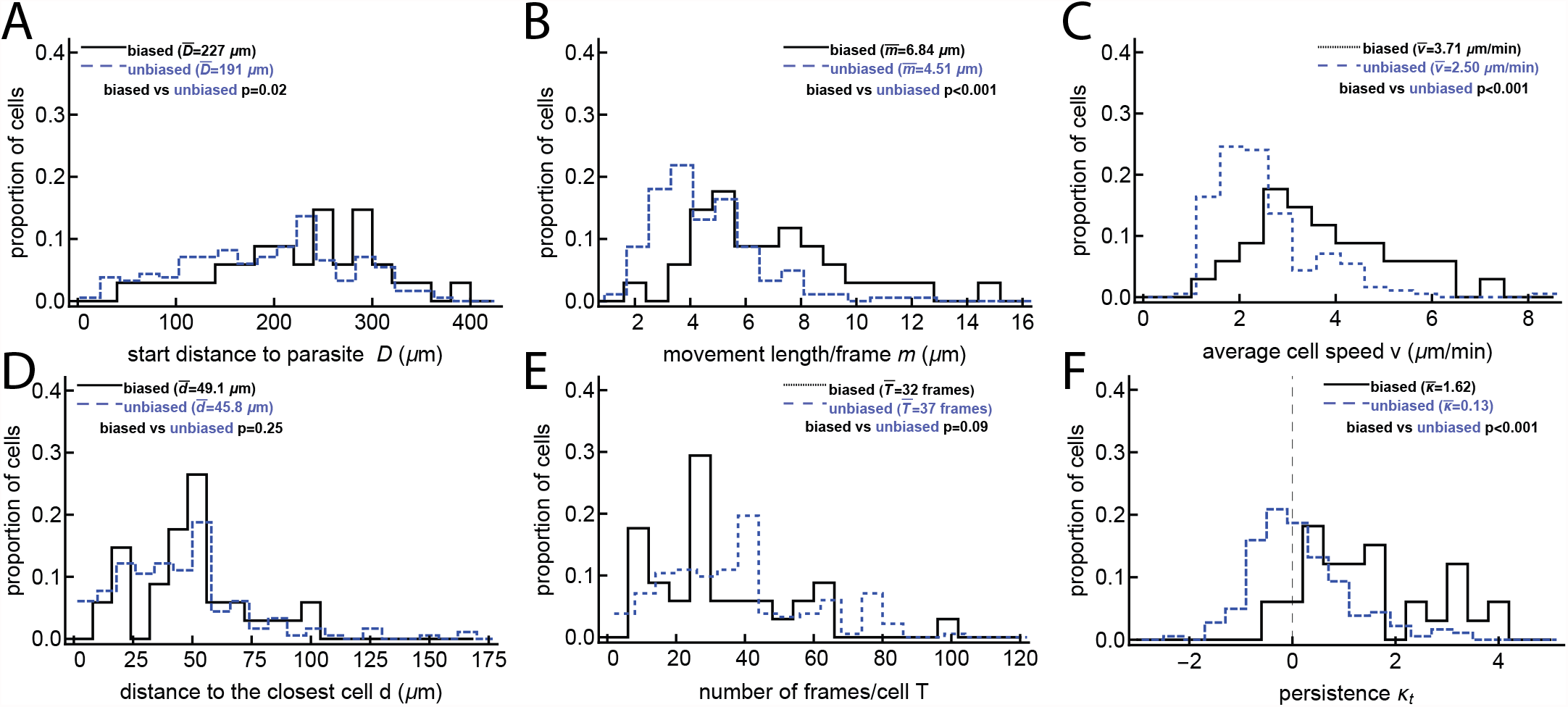
Speed and turning angle both correlate with detection of attraction when no or few T cells have found the parasite. These are metadata results from tests that use data divided into biased and unbiased T cells for the unclustered/small clustered dataset (dataset #1). For more details, see Figure 2.

**Figure S15:**
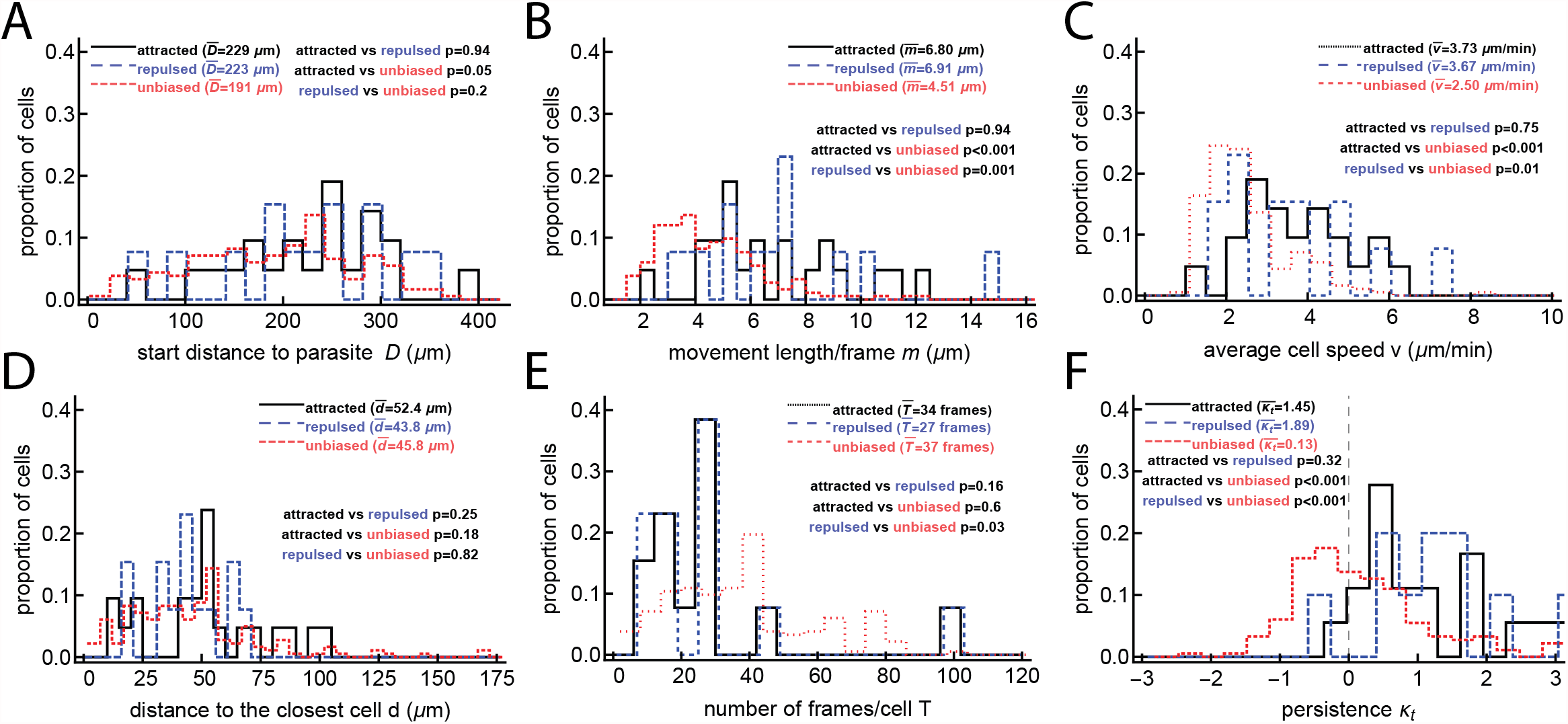
Speed and turning angle both correlate with detection of attraction when no or few T cells have found the parasite. These are metadata results from tests that use data divided into attracted, repulsed, and unbiased T cells for the unclustered/small clustered dataset. For more details, see Figure 2.

**Figure S16:**
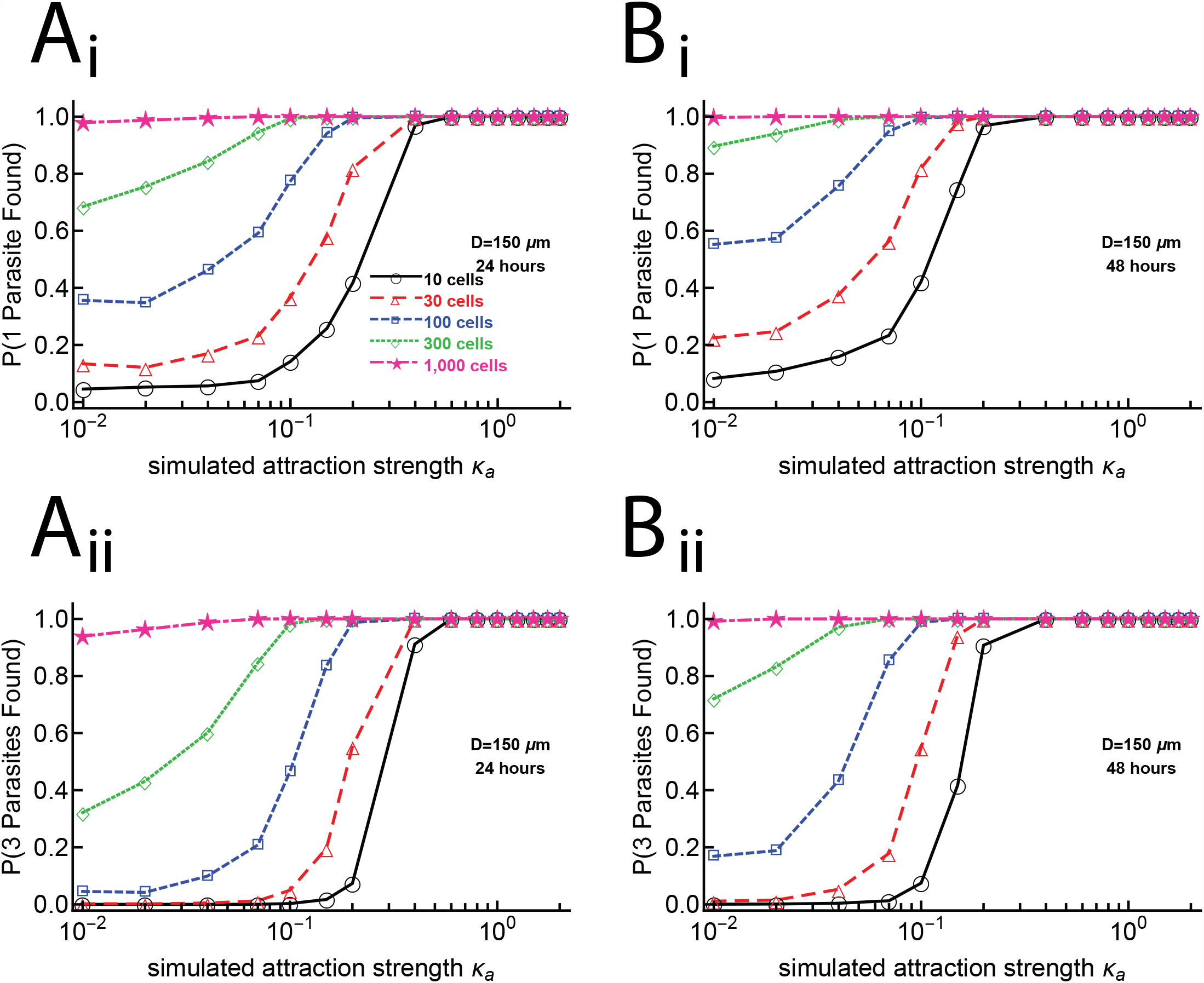
Many CD8 T cells per parasite are needed to ensure that all parasites are found within 48 hours after infection. Groups of 10, 30, 100, 300, and 1000 cells per parasite are simulated 1000 times for values of *κ*_*a*_ between 0 and 2 with a starting distances of 150 *μ*m and are “sampled” every two minutes for lengths of 24 (panels A) or 48 (panels B) hours. Each parasite that has a cell reach within 40*μ*m of it is considered a success. Panels i show the probability of a parasite being found, while panels ii show the probabilities of three parasites being found. If there are 100 cells per parasite, a *κ*_*a*_ of 1.0 is required for essentially all parasites to be found within 48 hours.

## References

1. https://www.who.int/malaria/publications/world-malaria-report-2018/report/en/.

2. Aleshnick, M., Ganusov, V. V., Nasir, G., Yenokyan, G. & Sinnis, P. Experimental determination of the force of malaria infection reveals a non-linear relationship to mosquito sporozoite loads. 16, e1008181. ISSN: 1553-7374. aheadofprint.

3. Murphy, J. R., Baqar, S., Davis, J. R., Herrington, D. A. & Clyde, D. F. Evidence for a 6.5-day minimum exoerythrocytic cycle for Plasmodi um falciparum in humans and confirmation that immunization with a synthetic peptide representative of a region of the circumsporozoite protein retards infection. J Clin Microbiol 27, 1434–1437 (1989).

4. Hermsen, C. C. et al. Detection of Plasmodium falciparum malaria parasites in vivo by real-time quantitative PCR. Mol Biochem Parasitol 118, 247–251 (2001).

5. Sturm, A. et al. Manipulation of host hepatocytes by the malaria parasite for delivery into liver sinusoids. Science 313, 1287–1290 (2006).

6. Miller, L. H., Ackerman, H. C., Su, X.-z. & Wellems, T. E. Malaria biology and disease pathogenesis: insights for new treatmen ts. Nat Med 19, 156–167 (2013).

7. Schmidt, N. W. et al. Memory CD8 T cell responses exceeding a large but definable threshold provide long-term immunity to malaria. Proc Natl Acad Sci U S A 105, 14017–22 (2008).

8. Schmidt, N. W., Butler, N. S., Badovinac, V. P. & Harty, J. T. Extreme CD8 T cell requirements for anti-malarial liver-stage immunity following immunization with radiation attenuated sporozoites. PLoS Pathog 6, e1000998 (2010).

9. Schmidt, N. W. & Harty, J. T. Cutting edge: attrition of Plasmodium-specific memory CD8 T cells results in decreased protection that is rescued by booster immunization. J Immunol 186, 3836–3840 (2011).

10. Cockburn, I. A. et al. In vivo imaging of CD8+ T cell-mediated elimination of malaria liver stages. eng. Proc Natl Acad Sci U S A 110, 9090–9095. http://dx.doi.org/10.1073/pnas.1303858110 (May 2013).

11. Kelemen, R. K., Rajakaruna, H., Cockburn, I. A. & Ganusov, V. V. Clustering of activated CD8 T cells around malaria-infected hepatocytes is rapid and is driven by antigen-specific cells. Frontiers in immunology 10, 2153. ISSN: 1664-3224 (2019).

12. Kelemen, R. K. et al. Classification of T Cell Movement Tracks Allows for Prediction of Cell Function. International Journal of Computational Biology and Drug Design (2014).

13. Krummel, M. F., Bartumeus, F. & Gérard, A. T cell migration, search strategies and mechanisms. Nature Reviews Immunology 16, 193–201. ISSN: 1474-1733 (2016).

14. Okada, T. et al. Antigen-Engaged B Cells Undergo Chemotaxis toward the T Zone and Form Motile Conjugates with Helper T Cells (B Cell-T Cell Interaction Dynamics). eng. PLoS Biology 3, e150. ISSN: 1544-9173 (2005).

15. Beltman, J. B., Allen, C. D. C., Cyster, J. G. & de Boer, R. J. B cells within germinal centers migrate preferentially from dark to light zone. eng. Proceedings of the National Academy of Sciences of the United States of America 108, 8755–8760. ISSN: 0027-8424 (2011).

16. Ariotti, S. et al. Subtle CXCR3-Dependent Chemotaxis of CTLs within Infected Tissue Allows Efficient Target Localization. eng. Journal of immunology (Baltimore, Md. : 1950) 195. ISSN: 1550-6606 (Dec. 2015).

17. Beltman, J. B., Marée, A. F. M. & de Boer, R. J. Analysing immune cell migration. Nat Rev Immunol 9, 789–798 (2009).

18. Turchin, P., Odendaal, F. J. & Rausher, M. D. Quantifying Insect Movement in the Field. Environ Entomol 20, 955–963. ISSN: 1938-2936 (1991).

19. Turchin, P. Quantitative Analysis of Movement: Measuring and Modeling Population Redistribution in Animals and Plants. 381. ISSN: 0033-5770 (1998).

20. Duchesne, T., Fortin, D. & Rivest, L.-P. Equivalence between Step Selection Functions and Biased Correlated Random Walks for Statistical Inference on Animal Movement. PloS one 10, e0122947. ISSN: 1932-6203 (4 2015). epublish.

21. Wang, L. T. et al. A Potent Anti-Malarial Human Monoclonal Antibody Targets Circumsporozoite Protein Minor Repeats and Neutralizes Sporozoites in the Liver. Immunity 53, 733–744.e8. ISSN: 1097-4180 (4 Oct. 2020). ppublish.

22. Cockburn, I. A., Tse, S.-W. & Zavala, F. CD8+ T cells eliminate liver-stage Plasmodium berghei parasites without detectable bystander effect. eng. Infect Immun 82, 1460–1464. http://dx.doi.org/10.1128/IAI.01500-13 (Apr. 2014).

23. McNamara, H. A. et al. Up-regulation of LFA-1 allows liver-resident memory T cells to patrol and remain in the hepatic sinusoids. Science Immunology, 1–10 (2 2017).

24. Hong, Y. On computing the distribution function for the Poisson binomial distribution. eng. Computational Statistics and Data Analysis 59, 41–51. ISSN: 0167-9473 (2013).

25. Mardia, K. V. Directional statistics Wiley; 2000.

26. Fisher, N. I. Statistical analysis of spherical data eng. Cambridge University Press, 1987.

27. Castellino, F. et al. Chemokines enhance immunity by guiding naive CD8+ T cells to sites of CD4+ T cell–dendritic cell interaction. Nature 440, 890–895. ISSN: 0028-0836 (2006).

28. Jakob, W. Numerically stable sampling of the von Mises Fisher distribution on S2 (and other tricks) http://www.mitsuba-renderer.org/~wenzel/files/vmf.pdf.

29. Jerison, E. R. & Quake, S. R. Heterogeneous T cell motility behaviors emerge from a coupling between speed and turning in vivo. eLife. https://elifesciences.org/articles/53933 (2020).

30. Fernandez-Ruiz, D. et al. Liver-Resident Memory CD8(+) T Cells Form a Front-Line Defense against Malaria Liver-Stage Infection. Immunity 45, 889–902. ISSN: 1097-4180 (4 Oct. 2016).

31. Marino, D. J. Age-specific absolute and relative organ weight distributions for B6C3F1 mice. Journal of toxicology and environmental health. Part A 75, 76–99. ISSN: 1528-7394 (2 2012). ppublish.

32. Olsen, T. M., Stone, B. C., Chuenchob, V. & Murphy, S. C. Prime-and-Trap Malaria Vaccination To Generate Protective CD8+ Liver-Resident Memory T Cells. J Immunol 201, 1984–1993. ISSN: 1550-6606 (7 Oct. 2018).

33. Gola, A. et al. Prime and target immunization protects against liver-stage malaria in mice. Science translational medicine 10. ISSN: 1946-6242 (460 Sept. 2018).

34. Krummel, M. F., Bartumeus, F. & Gérard, A. r. T cell migration, search strategies and mechanisms. Nature reviews. Immunology 16, 193–201. ISSN: 1474-1741 (Mar. 2016).

35. Okada, T. et al. Antigen-engaged B cells undergo chemotaxis toward the T zone and form motile conjugates with helper T cells. PLoS biology 3, e150. ISSN: 1545-7885 (June 2005).

36. Harris, T. H. et al. Generalized Lévy walks and the role of chemokines in migratio n of effector CD8+ T cells. Nature 486, 545–548 (June 2012).

37. Hoft, S. G. et al. The Rate of CD4 T Cell Entry into the Lungs during Mycobacterium tuberculosis Infection Is Determined by Partial and Opposing Effects of Multiple Chemokine Receptors. Infection and immunity 87. ISSN: 1098-5522 (6 June 2019).

38. Guidotti, L. G. et al. Immunosurveillance of the liver by intravascular effector CD8(+) T cells. Cell 161, 486–500. ISSN: 1097-4172 (3 Apr. 2015).

39. Rajakaruna, H., O’Connor, J., Cockburn, I. A. & Ganusov, V. V. Environment-imposed constraints make Brownian walkers efficient searchers BioRxiv. 2020. https://doi.org/10.1101/2020.11.06.371690.

40. Mrass, P. et al. ROCK regulates the intermittent mode of interstitial T cell migration in inflamed lungs. eng. Nat Commun 8, 1010–1010. ISSN: 2041-1723 (Dec. 2017).

41. Moses, M. E., Cannon, J. L., Gordon, D. M. & Forrest, S. Distributed Adaptive Search in T Cells: Lessons From Ants. Frontiers in Immunology 10, 1357. ISSN: 1664-3224 (2019). epublish.

## Supplemental references

42. Pawitan, Y. In All Likelihood: Statistical Modelling and Inference Using Likelihood 544 (Oxford University Press, 2001).

43. Ahmed, D. A. & Petrovskii, S. V. Analysing the impact of trap shape and movement behaviour of ground-dwelling arthropods on trap efficiency. Methods Ecol Evol 10, 1246–1264. ISSN: 2041-210X (2019).

44. Uhlenbeck, G. E. & Ornstein, L. S. On the Theory of the Brownian Motion. Phys. Rev. 36, 823–841. https://link.aps.org/doi/10.1103/PhysRev.36.823 (5 Sept. 1930).

45. Wu, P.-H., Giri, A., Sun, S. X. & Wirtz, D. Three-dimensional cell migration does not follow a random walk. Proceedings of the National Academy of Sciences 111, 3949–3954. ISSN: 0027-8424. eprint: https://www.pnas.org/content/111/11/3949.full.pdf. https://www.pnas.org/content/111/11/3949 (2014).

46. Vectors with a certain magnitude in Mathematica https://mathematica.stackexchange.com/questions/13038/vectors-with-a-certain-magnitude-in-mathematica/13042#13042.

